# Estimating genetic nurture with summary statistics of multi-generational genome-wide association studies

**DOI:** 10.1101/2020.10.06.328724

**Authors:** Yuchang Wu, Xiaoyuan Zhong, Yunong Lin, Zijie Zhao, Jiawen Chen, Boyan Zheng, James J. Li, Jason M. Fletcher, Qiongshi Lu

## Abstract

Marginal effect estimates in genome-wide association studies (GWAS) are mixtures of direct and indirect genetic effects. Existing methods to dissect these effects require family-based, individual-level genetic and phenotypic data with large samples, which is difficult to obtain in practice. Here, we propose a novel statistical framework to estimate direct and indirect genetic effects using summary statistics from GWAS conducted on own and offspring phenotypes. Applied to birth weight, our method showed nearly identical results with those obtained using individual-level data. We also decomposed direct and indirect genetic effects of educational attainment (EA), which showed distinct patterns of genetic correlations with 45 complex traits. The known genetic correlations between EA and higher height, lower BMI, less active smoking behavior, and better health outcomes were mostly explained by the indirect genetic component of EA. In contrast, the consistently identified genetic correlation of autism spectrum disorder (ASD) with higher EA resides in the direct genetic component. Polygenic transmission disequilibrium test showed a significant over-transmission of the direct component of EA from healthy parents to ASD probands. Taken together, we demonstrate that traditional GWAS approaches, in conjunction with offspring phenotypic data collection in existing cohorts, could greatly benefit studies on genetic nurture and shed important light on the interpretation of genetic associations for human complex traits.

## Introduction

Genome-wide association studies (GWAS) have been a great success in the past decade, identifying tens of thousands of associations for numerous complex human traits^1^. The standard GWAS approach estimates the marginal association between each single-nucleotide polymorphism (SNP) and a phenotype while assuming that genetic and environmental factors additively affect the phenotype. Despite the simplicity, such an analytical strategy is computationally efficient and statistically robust. However, interpretation of GWAS associations remains a challenge, in part because most identified associations have weak effect sizes and are located in the non-coding regions of the genome^2,3^. Interpretation is especially challenging for behavioral traits since the role of each variant or gene in complex human behavior is difficult to disentangle. Nevertheless, biobank-scale GWAS of complex traits have produced polygenic scores (PGS) that aggregate the effects of many SNPs in the genome to provide robust prediction of trait values^4^. These scores are widely used in social genomics research, although our understanding of the underlying mechanism is superficial and incomplete^5^.

Recent evidence from family-based studies suggested that a substantial fraction of genetic associations may be mediated by the family environment^6-16^. In particular, parental genotypes could affect the family environment through the parents’ educational attainment^17^, personalities^18,19^, behavior^20-24^, and socioeconomic status^25^, which could subsequently affect the offspring’s phenotypes^26^. As a result, a person’s genotypes, which also reside in his or her biological parents, could associate with the person’s phenotype both directly (through biological processes) and indirectly (through parents and the family environment they create). Due to the correlation between parental and offspring genotypes, GWAS captures both the direct and indirect genetic effects in its estimates, which further complicates the interpretation of GWAS results^13^. If the genetic nurture effect (i.e., parental genotypes affecting offspring phenotype) is present for a given trait, downstream analyses based on GWAS associations could be biased and misleading^6,8,27^.

It is thus crucial to decompose the direct and indirect genetic effects and understand how they jointly affect the phenotype. By leveraging large-scale trio cohorts and regressing the offspring phenotype on two sets of PGS calculated using transmitted and non-transmitted alleles in parents, Kong et al.^6^ convincingly demonstrated the existence of genetic nurture effects for multiple traits. In particular, PGS of non-transmitted alleles in parents has an effect size that is about 30% of that by the standard PGS for educational attainment (EA). Using PGS, several other studies^7-12^ also identified indirect genetic effects on various phenotypes. Existing methods to detect direct and indirect genetic effects, however, have limitations. First, they require individual-level genotype and phenotype data of a large number of parents-offspring trios, or in some cases, other types of rare samples (e.g., adopted individuals^11,12^). Although sample size in GWAS has been steadily increasing, number of trio samples with accessible individual-level data remains moderate even in large biobanks. Second, existing methods quantify genetic nurture using PGS which relies on large GWAS conducted on samples independent from the study. Even when such a GWAS exists, it remains challenging to interrogate the direct and indirect effects of each SNP using designs and data similar to the current GWAS practice, which is critical for functional follow-ups and out-of-sample prediction^13^.

Although a simple study design that regresses the phenotypes on both own and parental genotypes should provide estimates for direct and indirect genetic effects of each SNP, such a strategy is most likely underpowered given the limited sample size of trios in existing cohorts. Several recent studies have attempted to solve this challenging problem. Warrington et al.^28^ used a structural equation model (SEM) approach to decompose direct genetic effects and indirect maternal effects on birth weight while assuming paternal effects to be 0. This approach only requires summary statistics from a standard GWAS on birth weight and a second GWAS based on maternal genotypes and offspring phenotypes, thus effectively expanding the available sample size. However, the SEM approach was too computationally demanding to be applied to the genome-wide scale and a “weighted linear model” alternative could not account for sample overlap if individual-level data are unavailable. Another recent approach^14,15^ expands family genotype data by imputing the unobserved parental genotypes using data from other family members. However, this approach still requires a large sample of sibling or parent-offspring pairs. Further, when parental genotypes are imputed from sibling pairs, it is challenging to distinguish paternal and maternal autosomal genotypes. Thus, separate estimation of indirect maternal and paternal effects is unattainable.

Here, we introduce DONUTS (**d**ecomp**o**sing **n**ature and n**u**r**t**ure using GWAS **s**ummary statistics), a novel statistical framework that can estimate direct and indirect genetic effects at the SNP level. It requires GWAS summary statistics as input, allows differential paternal and maternal effects, and accounts for GWAS sample overlap and assortative mating. DONUTS has low computational burden and can complete genome-wide analyses within seconds. Applied to birth weight, our method showed near-identical effect estimates compared to analyses^28^ that leveraged individual-level data and improved standard error and statistical power after accounting for sample overlap. We also applied our method to dissect the direct and indirect genetic effects of EA. Our results revealed distinct genetic correlations of the direct and indirect genetic components of EA with various traits and shed important light on the complex and heterogenous genetic architecture of EA. Followed up in three independent cohorts of ASD proband-parent trios, we identified significant over-transmission of the direct component of EA from healthy parents to ASD probands but not to the healthy siblings.

## Results

### Overview of the methods

The key idea of our statistical framework is illustrated in **Figure 1**. Derivations and statistical details are shown in **Methods** and **Supplementary Note**. If genetic data are available in a number of parents-offspring trios, by regressing the offspring phenotype values *Y*_*M*_ on the offspring, maternal, and paternal genotypes (i.e., *G*_O_, *G*_M_, and *G*_P_) for a given SNP, the coefficients in the joint regression represent the direct genetic effect *β*_dir_, indirect maternal effect *β*_ind_mt_, and paternal effect *β*_ind_pt_, respectively. We could write this model as

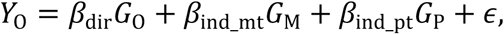

where *ϵ* is the environmental noise. The total contribution of parental genotypes on offspring phenotype, (*β*_ind_mt_*G*_M_ + *β*_ind_pt_*G*_P_), can be further partitioned into the contribution of transmitted alleles (*β*_ind_mt_*T*_M_ + *β*_ind_pt_*T*_P_) and non-transmitted alleles (*β*_ind_mt_*NT*_M_ + *β*_ind_pt_*NT*_P_). In our framework, we define the indirect genetic effect *β*_ind_ as the effect of a person’s genotype on the phenotype via the indirect pathway that goes through biological parents and the family environment. The component of parental indirect contribution that can be affected by *G*_O_ is (*β*_ind_mt_*T*_M_ + *β*_ind_pt_*T*_P_). Regressing it on *G*_O_, we can obtain the indirect genetic effect *β*_ind_ = (*β*_ind_mt_ + *β*_ind_pt_)/2. Unsurprisingly, the indirect effect size is the average of the indirect maternal and paternal effects since each parent contributes half of the offspring’s genotype. A key question we aim to investigate in this paper is whether it is possible to estimate the direct and indirect effect sizes (i.e., *β*_dir_, *β*_ind_, *β*_ind_mt_, and *β*_ind_pt_) from marginal GWAS association statistics via proper study designs.

**Figure 1.**
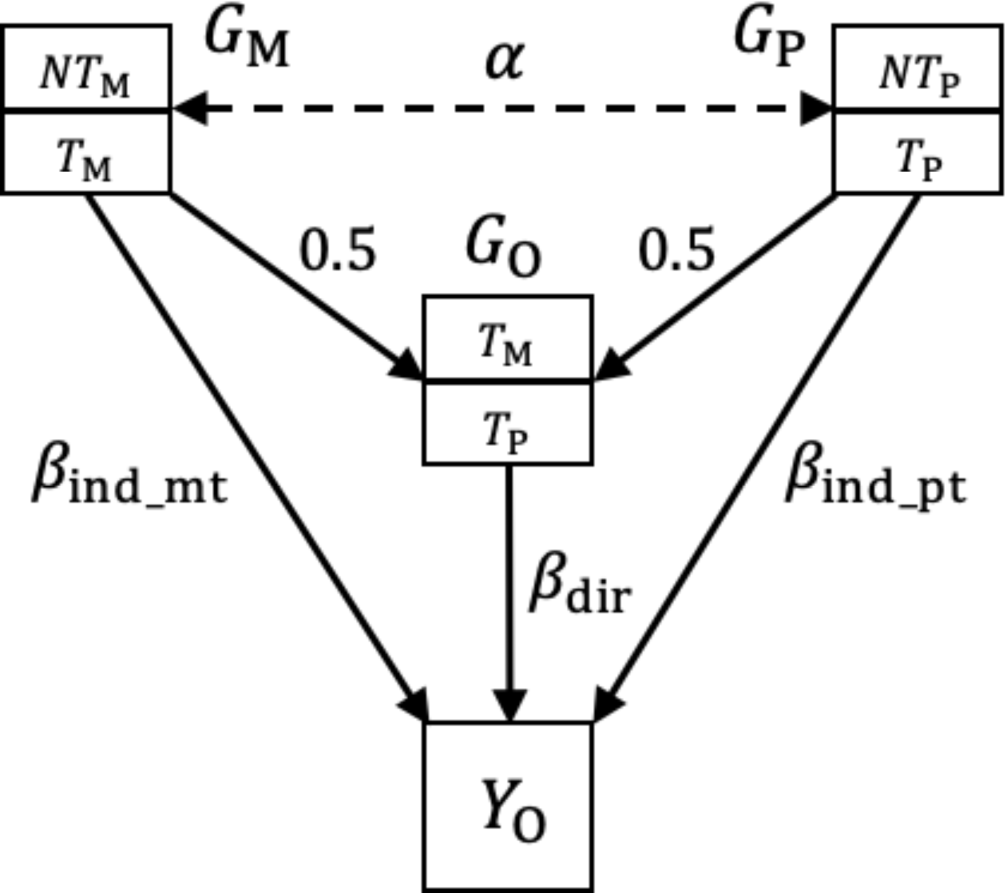
Schematic diagram of direct and indirect genetic effects. *G*_*M,P,O*_ represents the maternal, paternal, and offspring genotypes, respectively. *α* = Corr(*G*_*M*_, *G*_*P*_) is the correlation between spousal genotypes at a locus. Effect size 0.5 is due to the fact that half of the parent’s genome is randomly transmitted to the offspring *G*_*O*_. *Y*_*O*_ is the offspring’s phenotype. *T* and *NT* represent transmitted and non-transmitted alleles from a parent to the offspring. In general, both offspring and parental genotypes could affect the offspring’s phenotype with effect sizes of *β*_dir_, *β*_ind_mt_ and *β*_ind_pt_, respectively.

Instead of focusing on a joint regression based on trio data, we describe three separate GWAS. We refer to the marginal regressions of own phenotype (*Y*_O_) on own genotypes (*G*_O_) as GWAS-O. GWAS-M and GWAS-P denote the marginal analyses that regress offspring phenotype on maternal and paternal genotypes (i.e., *G*_M_ and *G*_P_), respectively. *β*_O_, *β*_M_, and *β*_P_ denote the expectation of marginal effect estimates obtained from these three analyses. It can be shown that *β*_dir_ and *β*_ind_ of a given SNP are linear combinations of *β*_O_, *β*_M_, and *β*_P_ (**Methods** and **Supplementary Note**):

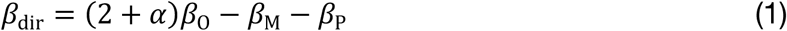

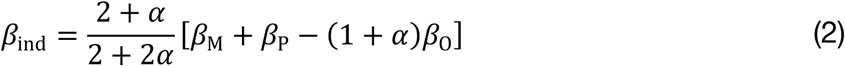

where *α* = Corr(*G*_*M*_, *G*_*P*_) is the correlation between spousal genotypes at the locus, which quantifies the degree of assortative mating. Plugging in the ordinary least squares estimates 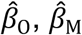, and 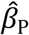 from the three marginal GWASs described above, we obtain the unbiased estimates for the direct and indirect effects of each SNP. Importantly, we do not require *G*_O_, *G*_M_ and *G*_P_ to be obtained from actual trios. In fact, samples in the three GWAS could be independent or partially overlapped. From the equations above, we also found that

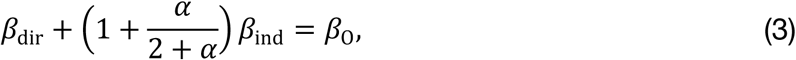

which clearly shows that the effect size from a typical GWAS is the combination of both direct and indirect effects and is also affected by assortative mating.^13^

Besides direct and indirect effects (i.e., *β*_dir_ and *β*_ind_), we could also derive the expressions for indirect maternal and paternal effects (i.e., *β*_ind_mt_ and *β*_ind_pt_), which makes it possible to infer the parent-of-origin of genetic nurture. The results are summarized in **Table 1**. Case (i) is the most general scenario, where we use summary statistics from GWAS-O, GWAS-M, and GWAS-P to estimate *β*_dir_, *β*_ind_, *β*_ind_mt_, and *β*_ind_pt_. Case (ii) illustrates that it is not always necessary to have separate paternal and maternal GWASs. If paternal and maternal effects are identical (*β*_ind_mt_ = *β*_int_pt_) or if there are equal numbers of mothers and fathers (*n*_M_ = *n*_P_) in a parental GWAS (referred to as GWAS-MP where fathers and mothers from different families are pooled together in the GWAS), the corresponding effect size *β*_MP_ = (*n*_M_*β*_M_ + *n*_P_*β*_P_)/(*n*_M_ + *n*_P_) can be used to estimate *β*_dir_ and *β*_ind_. Case (iii) illustrates a special case where only maternal genotype has an indirect effect while the paternal effect is zero (**Supplementary Figure 1A**). If we further assume random mating (*α* = 0), then our model gives identical estimates for direct effect *β*_dir_ and maternal effect *β*_ind_mt_ compared to previous work on birth weight^28^. The results for the case with only indirect paternal effects are similar.

**Table 1.**
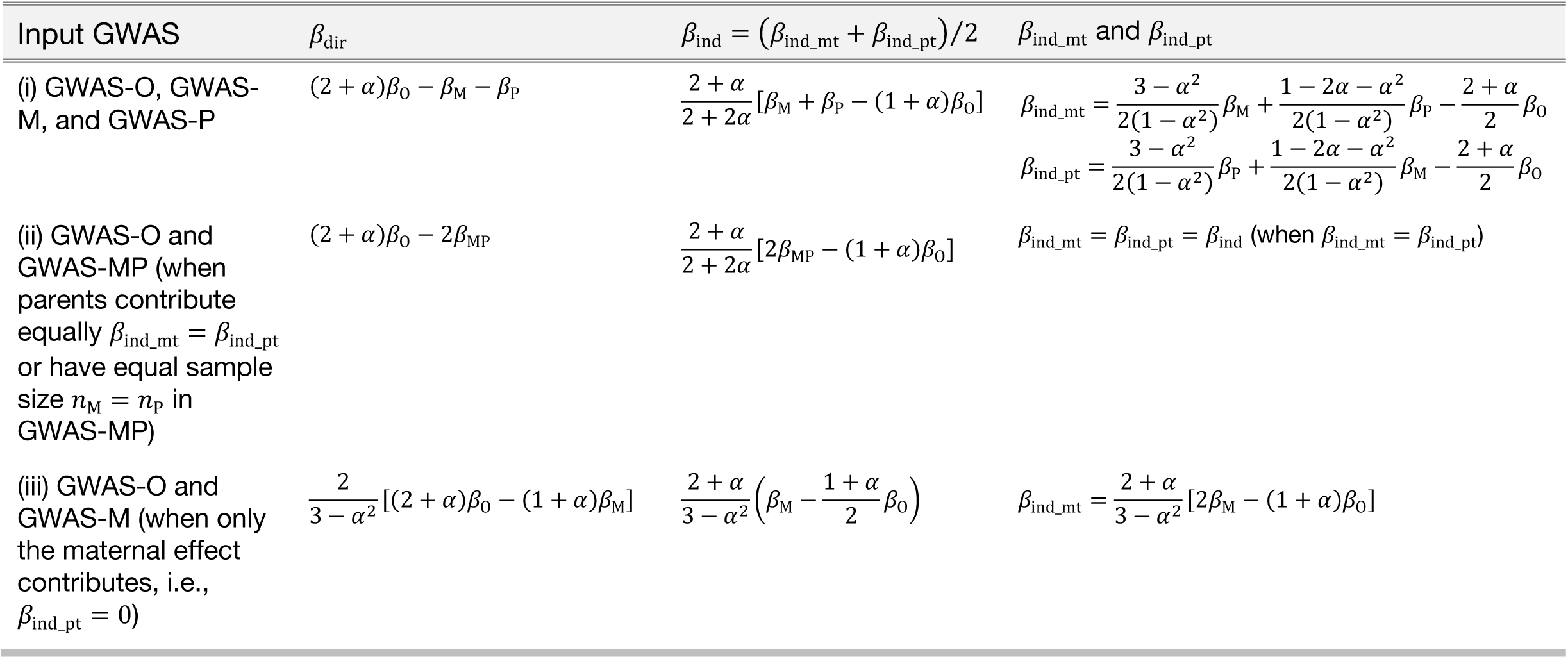
Estimating direct and indirect genetic effects from multi-generational GWAS summary statistics. We illustrate the direct and indirect effect sizes under three different settings. (i) is the general case where GWAS-O, GWAS-M, and GWAS-P are used as input. In case (ii), GWAS-O and GWAS-MP are used. This is valid only when *β*_ind_mt_ = *β*_ind_pt_ or *n*_M_ = *n*_P_. If we only know *n*_M_ = *n*_P_, we cannot obtain separate estimates for the indirect maternal and paternal effects. Case (iii) is when the indirect paternal effect size is 0. *β*_O, M, P, MP_ are the expected effect sizes in GWAS-O, GWAS-M, GWAS-P, and GWAS-MP, respectively. In all the cases, we always have *β*_dir_ + [1 + *α*/(2 + *α*)]*β*_ind_ = *β*_O_ and *β*_ind_ = 5*β*_ind_mt_ + *β*_ind_pt_)/2.

Calculations of the variances of estimated direct and indirect effects are straightforward when the input GWASs are independent. However, it is possible for a subset of individuals to be involved in both the GWAS of their own phenotype and the GWAS of their children’s phenotype (**Supplementary Figure 1B**), which causes technical correlations among 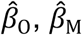, and 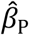. We show that the correlations can be estimated using the intercept term from linkage disequilibrium score (LDSC) regression^29^ (**Supplementary Note**), thereby correcting the sample overlap bias in standard error estimates.

### Simulation results

We performed extensive simulations to demonstrate that our method provides unbiased estimates for direct and indirect effects, shows well-controlled type-I error, and properly accounts for sample overlap (**Methods** and **Supplementary Figure 2**). The results are summarized in **Figure 2** and **Supplementary Figures 3-5. Figures 2A** and **2C** describe results for case (i) in **Table 1** where three sets of GWAS summary statistics are used. The estimates for direct, indirect, indirect maternal, and indirect paternal effect sizes were all unbiased. When only GWAS-O and GWAS-MP are available (case ii in **Table 1**), we could not distinguish indirect maternal and paternal effects but could still estimate the indirect genetic effect (**Figure 2B**). Here, despite the difference between indirect maternal and paternal effect sizes, estimation of the indirect genetic effects remained unbiased when equal number of fathers and mothers were used in GWAS-MP.

**Figure 2.**
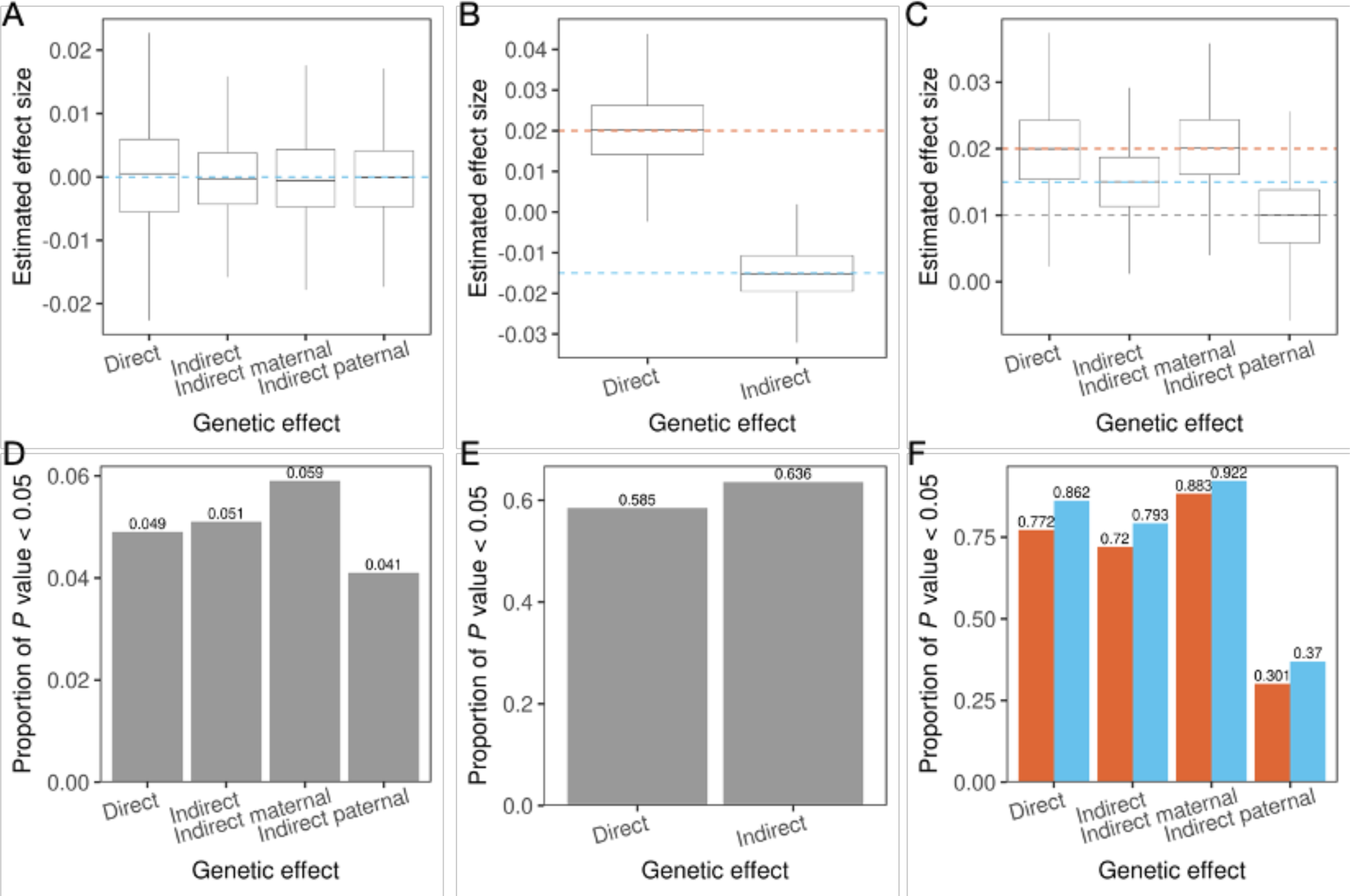
Simulation results. Box plots of direct and indirect effect size estimates (**A-C**) and the proportion (numbers shown at the top of each bar) of *p* values smaller than 0.05 (**D-F**). Each column shows results for the same simulation setting. Red and blue dashed lines indicate true values of direct and indirect genetic effects and grey dashed lines are the true indirect maternal and indirect paternal effect sizes. The direct, indirect maternal, and indirect paternal effect sizes are (0, 0, 0), (0.02, -0.02, -0.01), and (0.02, 0.02, 0.01) for panels **A**-**C**, respectively. Panels **A** and **C** describe results for case (i) in **Table 1** where three input GWAS are used. Panel **B** describes case (ii) where GWAS-O and GWAS-MP are used as input. There are no sample overlaps in **A** and **B** and a complete overlap in **C**, i.e., all samples in GWAS-M and GWAS-P are also in GWAS-O. In **F**, blue and red bars show the statistical power with and without sample overlap correction, respectively. *n*_O_ = *n*_M_ = *n*_P_ = 30K in **A** and **B**. *n*_O_ = *n*_M_ + *n*_P_ = 60K, *n*_M_ = *n*_P_ = 30K in **C**.

Sample overlap in input GWASs will not affect effect size estimation. However, it will affect their standard errors due to the introduced correlations among 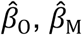, and 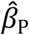. In **Figures 2C** and **2F**, there were overlapping samples between GWAS-O and parental GWAS. Since the phenotypic correlation among the overlapping samples (i.e., correlation between parental and offspring phenotypes) would most likely be positive, the covariance between effect size estimates is positive. As a result, correcting for sample overlap reduces standard error and increases power. Simulations under diverse settings all showed consistent results (**Supplementary Figures 3-5**).

### Direct and maternal effects on birth weight

To assess the performance of our framework, we applied DONUTS to dissect the direct genetic effect and maternal genetic effect on birth weight. Following a previous study^28^, we assumed random mating and absent paternal effect on offspring birth weight, which reduces the problem to a special case in our framework (case iii in **Table 1**; **Supplementary Figure 1**). Using summary statistics from GWAS-O and GWAS-M (N = 298,142 and 210,267, respectively; **Methods**), we estimated the direct and maternal effects of each SNP. Both estimates were highly concordant with the previous reports (Pearson correlations = 0.976 and 0.982, respectively; **Supplementary Figure 6**). The genetic correlations among these effects were very close to those reported in previous work (**Supplementary Tables 1** and **2**).

Of note, the UK Biobank (UKB)^30^ was a main cohort used in both GWAS-O and GWAS-M of birth weight, which caused a substantial sample overlap between two analyses. Warrington et al.^28^ addressed this problem by creating two linearly-transformed, orthogonal phenotypes for each individual who reported both her own birth weight and her first child’s birth weight. GWAS were then performed on the two new phenotypes. This approach requires individual-level genotype and phenotype data and thus is not easily applicable to other studies where only summary statistics are available. In fact, due to limited access to non-UKB samples, a small proportion of overlapping samples in the input GWAS were not accounted for in their study. Therefore, compared with our results, the standard errors given by the paper showed a mild inflation (**Supplementary Figure 6**).

To further demonstrate that our method could effectively account for sample overlap, we conducted GWAS-O and GWAS-M using 75,711 independent female samples of European ancestry in the UKB who reported birth weight of themselves and of their oldest child (**Methods**). Using these two sets of summary statistics with a complete sample overlap, we estimated the direct and indirect maternal effects of each SNP. For comparison (**Figure 3**), we followed Warrington et al.^28^ to run two separate GWAS on the orthogonal phenotypes representing the direct and maternal components of birth weight constructed using individual-level data. Results from these two approaches were nearly identical (Pearson correlation = 1.00 for both the direct and indirect effect estimates). Not properly accounting for sample overlap did not affect the effect size estimates but substantially inflated standard errors which led to reduced statistical power (**Figure 3**).

**Figure 3.**
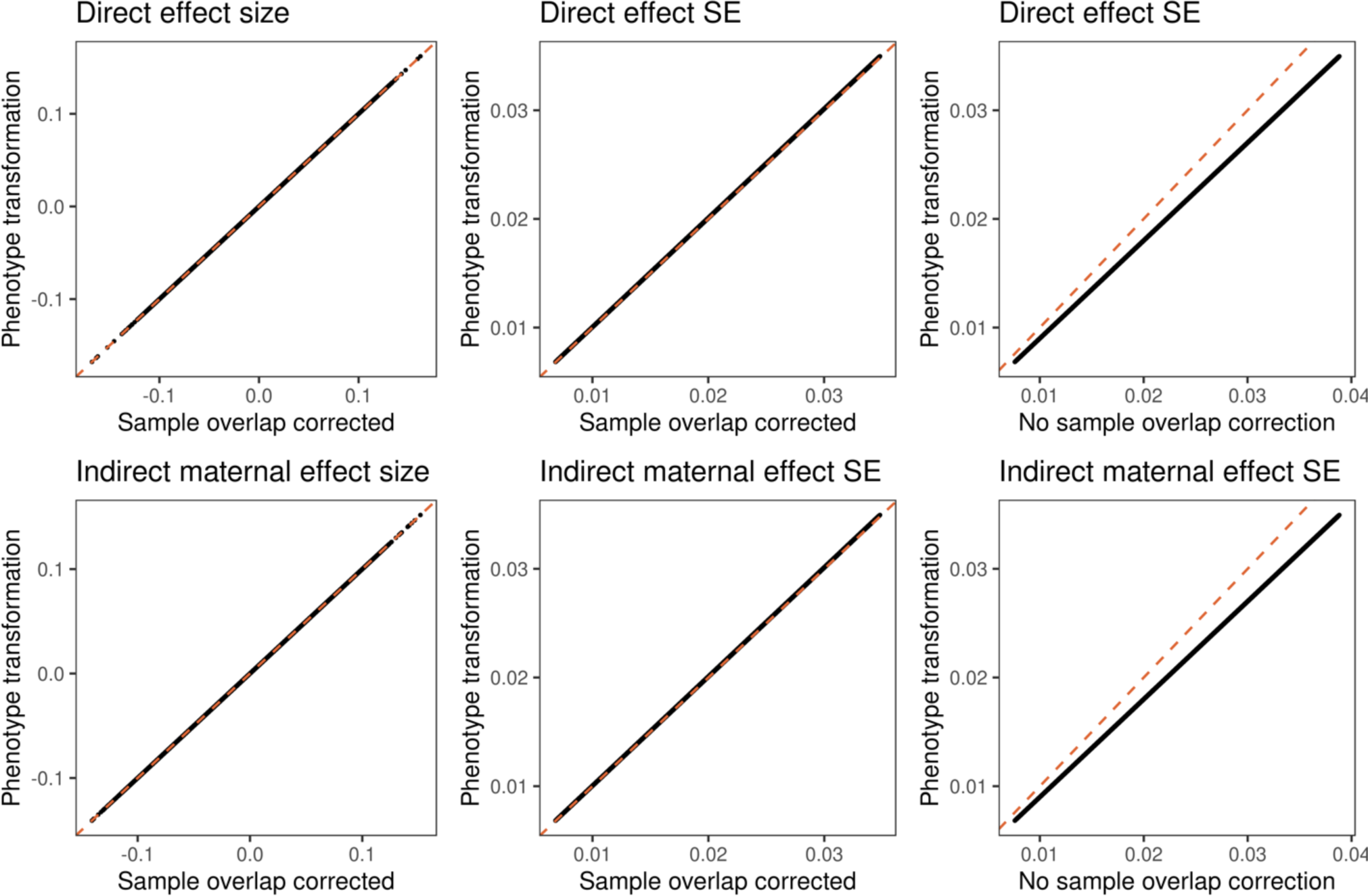
Comparison of DONUTS and analyses based on individual-level data. We estimated direct and indirect maternal genetic effects on birth weight using independent female samples of European ancestry in the UKB who reported both their own birth weight and their first child’s birth weight (N = 75,711). The x-axis of each panel shows results based on DONUTS and the y-axis shows results based on the phenotype transformation approach. Each data point represents a SNP. The 1^st^ row is for the direct genetic effect and the 2^nd^ row is for the indirect maternal effect. Two methods gave almost identical estimates for effect sizes (1^st^ column) and standard errors (2^nd^ column). The standard errors (SE) showed inflation if sample overlap was not accounted for (3^rd^ column). The diagonal line is highlighted in each panel.

### Partitioning direct and indirect genetic effects on educational attainment

Next, we conducted a GWAS on offspring EA using a total of 15,277 individuals from the UKB, Wisconsin Longitudinal Study (WLS), and Health and Retirement Study (HRS) while adjusting for year of birth, sex, genetic principal components (PCs), and cohort specific covariates (**Methods**). Due to the limited sample size, balanced sex ratio, and previous reports on comparable maternal and paternal effects on EA^6^, we pooled fathers and mothers together to perform a parental GWAS (i.e., GWAS-MP). Combining results in GWAS-MP with a meta analyzed GWAS-O that does not contain full sibling pairs in the UKB (N = 680,881), we estimated the direct and indirect effects on EA. Further, we applied SNIPar^15^ to impute the parental genotypes of full sibling pairs in the UKB and estimated direct and indirect effects with linear mixed models (**Methods**). We meta-analyzed two sets of analyses to obtain the final partitioned direct and indirect genetic effects on EA (effective N = 24,434 and 37,081 for direct and indirect effects, respectively). The flowchart of the analysis is illustrated in **Supplementary Figure 7**. No loci reached genome-wide significance at the current sample size (**Supplementary Figure 8**). We assumed random mating in the main analysis, but the results were highly robust to assortative mating (**Supplementary Note**; **Supplementary Figure 9**).

We estimated genetic correlations of the direct and indirect EA effects with 45 other complex traits using LDSC^29^ (**Figure 4** and **Supplementary Tables 3-6**). As a comparison, an alternative approach (i.e., GNOVA^31^) also showed consistent results (**Supplementary Figure 10** and **Supplementary Table 7**). At a false discovery rate (FDR) cutoff of 0.05, we identified 18 significant genetic correlations, 4 of which were with the direct effect and 14 were with the indirect effect, which highlighted the substantial contribution of genetic nurture on the etiologic sharing among complex traits. We also estimated genetic correlations based on a standard EA GWAS (i.e., GWAS-O; **Supplementary Figure 11**).

**Figure 4.**
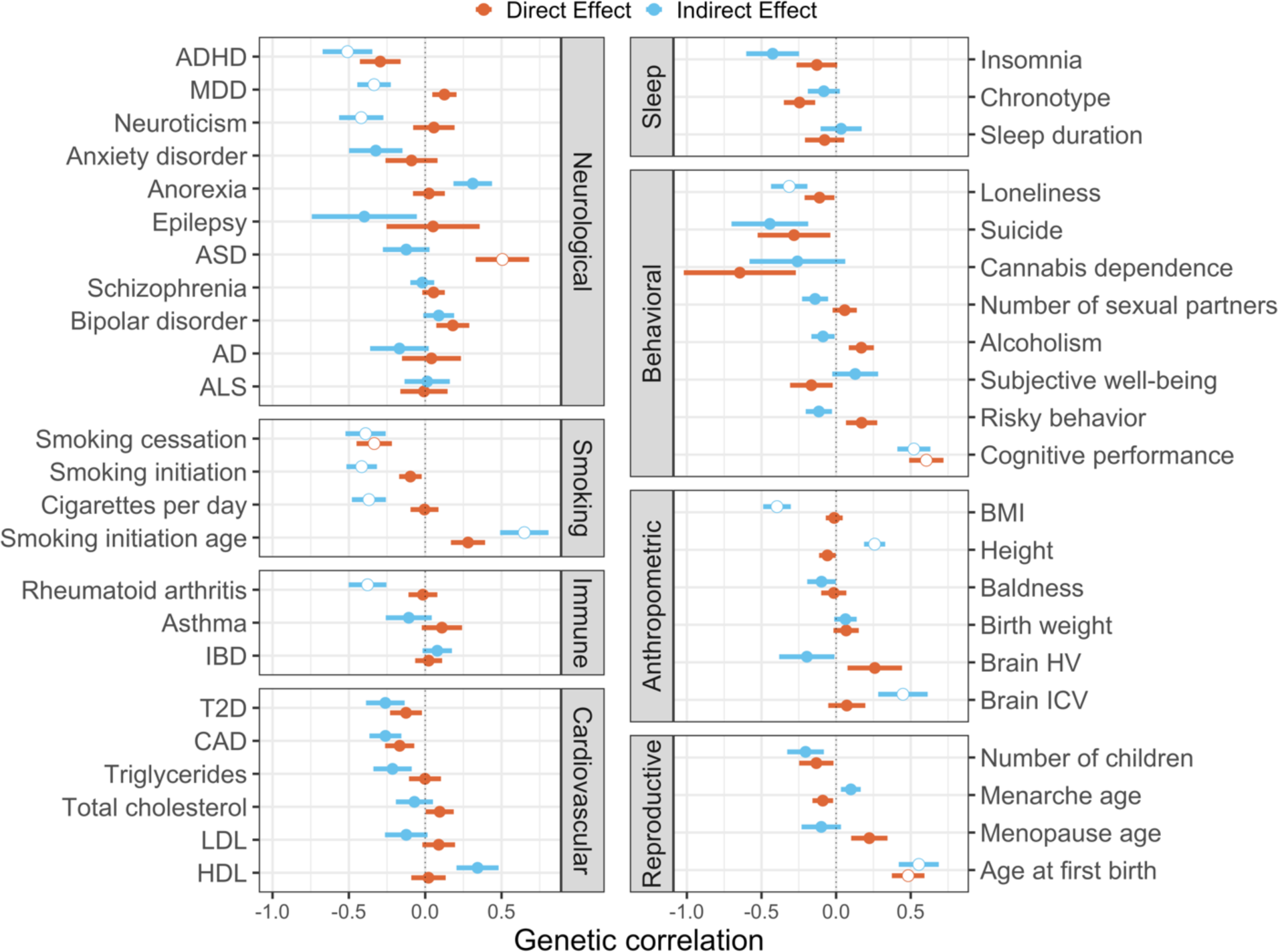
Genetic correlations of EA (direct and indirect effects) with 45 complex traits. Dots and intervals indicate the point estimates and standard error of genetic correlations, respectively. Significant correlations at an FDR cutoff of 0.05 are highlighted with white circles. ADHD: attention deficit/hyperactivity disorder; MDD: major depressive disorder; ASD: autism spectrum disorder; AD: Alzheimer’s diseases; ALS: amyotrophic lateral sclerosis; IBD: inflammatory bowel disease; T2D: type-2 diabetes; CAD: coronary artery disease; LDL and HDL: low and high-density lipoprotein; BMI: body-mass index; HV: hippocampal volume; ICV: intracranial volume.

Three traits, i.e., cognitive performance (*p* = 1.53 × 10^−7^ and 2.51 × 10^−6^), age at first birth (*p* = 1.02 × 10^−5^ and 3.64 × 10^−5^), and smoking cessation (*p* = 3.97 × 10^−3^ and 3.05 × 10^−3^), were significantly correlated with both direct and indirect components of EA. Across four traits for smoking behavior, we observed a consistent pattern that higher EA, especially its indirect component, was correlated with reduced smoking activity. Among neurological traits, attention-deficit/hyperactivity disorder (ADHD; *p* = 1.77 × 10^−3^), major depressive disorder (MDD; *p* = 2.27 × 10^−3^), and neuroticism (*p* = 3.87 × 10^−3^) showed significant negative correlations with the indirect EA effect while autism spectrum disorder (ASD; *p* = 3.91 × 10^−3^) was positively correlated with the direct effect. Notably, several diseases and anthropometric traits known to genetically correlate with EA, e.g., rheumatoid arthritis (*p* = 2.23 × 10^−3^), height (*p* = 2.77 × 10^−4^), and body-mass index (BMI; *p* = 1.85 × 10^−5^), were only correlated with the indirect component of EA in our analysis. Such a pattern was also observed for type-2 diabetes (T2D), coronary artery disease (CAD), and various lipid traits despite not reaching statistical significance.

Next, we assessed the predictive performance of PGS of direct and indirect effects on EA. We generated bioinformatically fine-tuned PGS^32^ for direct and indirect components of EA using UKB participants (**Methods; Supplementary Table 8**). Overlapping UKB samples were removed from the input GWAS when necessary (**Methods**). **Supplementary Figure 12** shows the predictive performance on 15,580 full sibling pairs and 370,308 independent UKB samples. Both direct and indirect PGS were significantly associated with EA in independent samples (*p* = 4.63 × 10^−8^ and 1.46 × 10^−9^) with similar effect sizes (regression coefficient = 8.7 × 10^−3^ and 9.6 × 10^−3^). Direct effect PGS was positively associated with the EA in full sibling pairs with an effect size comparable to that in the population (regression coefficient = 0.013). The indirect PGS was negatively correlated with EA in full siblings. However, due to a limited sample size, neither direct nor indirect PGS reached statistical significance in sibling pairs (p = 0.16 and 0.52). The effect sizes of these PGS were also substantially weaker compared to the standard EA PGS based on population GWAS^17^.

We found that the somewhat surprising yet consistently replicated genetic correlation between ASD and higher EA^33,34^ is mainly driven by the direct genetic component of EA (**Figure 4**). We followed up this finding in 7,804 ASD proband-parent trios from three cohorts (**Methods**), including the Autism Genome Project (AGP), Simons Simplex Collection (SSC), and Simons Foundation Powering Autism Research for Knowledge (SPARK). We performed polygenic transmission disequilibrium test^35^ (pTDT) to quantify the deviation of ASD probands’ EA PGS from the parents’ PGS (**Methods**). We identified a significant (*p* = 1.25 × 10^−3^) over-transmission of the direct effect EA PGS from healthy parents to ASD probands (**Figure 5** and **Supplementary Table 9**). We did not identify a significant over-transmission of the indirect EA PGS (*p* = 0.61). Neither PGS showed any significant deviation from transmission equilibrium in healthy sibling controls.

**Figure 5.**
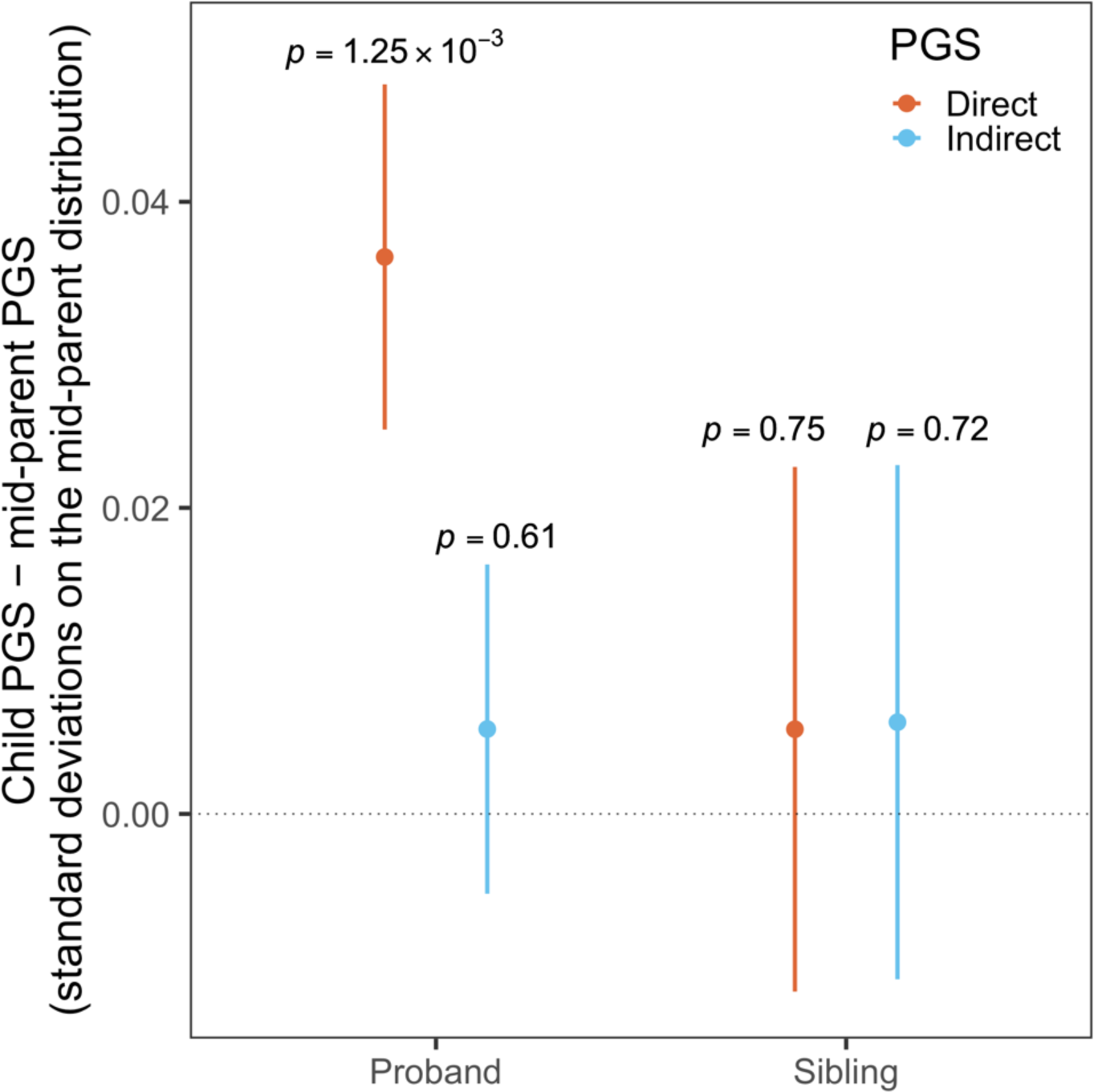
pTDT results for direct and indirect EA PGS in 7,804 ASD probands and 3,242 healthy siblings. Dots are the mean difference between child PGS and mid-parent PGS and intervals indicate the standard error.

## Discussion

GWAS has identified more than 60,000 genetic associations for thousands of human diseases and traits, yet our understanding towards their etiology remains incomplete^36^. Recent advances in family-based studies^6-9,14,15,28,37^ have convincingly demonstrated genetic nurture effects on a variety of behavioral traits as well as health-related outcomes. These results also shed important light on the limitations of current GWAS approaches. Accurate dissection of direct and indirect genetic effects is critical for advancing the interpretation of genetic associations and may fundamentally change the current practice of genetic prediction and its clinical applications.

In this paper, we introduced a novel statistical framework that uses summary statistics from multi-generational GWAS to decompose the direct and indirect genetic effects for a given trait. Compared to existing methods, our approach does not require access to individual-level data, has minimal computational burden, and accounts for GWAS sample overlap and assortative mating. In addition, when results from GWAS-M and GWAS-P are available, our method can partition the contribution of maternal and paternal genetic effects, thereby inferring the parent-of-origin of genetic nurture. Even when only a combined parental GWAS (i.e., GWAS-MP) is available, statistical inference of direct and indirect effects remain valid under weak assumptions. Importantly, due to these methodological advances, our approach does not require drastic changes to the current GWAS practice. All it needs is collecting offspring phenotype data (but not genotypes) in GWAS cohorts, which is substantially more economical and practical compared to collecting both genotypes and phenotypes from a large number of families. We note that even when individual-level data are available, our method will not have substantially lower power, especially for traits with higher heritability. We compared the effective sample sizes between our study design and a trio-based design (**Supplementary Note**) and found that the effective sample size of two approaches converge (**Supplementary Figure 13**).

EA is an important and highly complex trait that correlate with many health and social outcomes^17^. Kong et al.^6^ quantitatively demonstrated the existence of indirect genetic effects on EA. It is thus of great interest to understand the etiologic relevance of its direct and indirect components and how they affect other genetically correlated phenotypes. Using a PGS approach, Willoughby et al.^9^ found that the indirect effect of EA may work through the family socioeconomic status. The genetic relationships of the direct and indirect effect of EA with other traits, however, are still unknown. We dissected the genetic effects of EA at the SNP level using our approach. The direct and indirect components of EA showed distinct genetic correlations with other complex traits. The known genetic correlations between EA and higher height, lower BMI, less active smoking behavior, and better health outcomes were mostly explained by the indirect genetic component of EA, suggesting that parents with these traits may show stronger nurture effects on their children’s EA. One exception that stood out in our analysis was ASD, a clinically heterogenous neurodevelopmental disorder that has been consistently identified to genetically correlate with higher cognitive ability^33,34^. We found that the positive ASD-EA genetic correlation mostly resides in the direct component of EA. Followed up in three independent cohorts of ASD proband-parent trios, we identified significant over-transmission of the direct component of EA from healthy parents to ASD probands but not to the healthy siblings. These results added value to the recent advances in understanding common genetic variations’ roles in ASD etiology^33,35,38,39^ and provided critical new insights into the shared genetic basis of ASD and cognitive ability. These results also call for extreme caution in human genome editing and embryo screening^40^. Beyond the ethical issues, elevating the PGS of EA may have limited direct protective effects on health outcomes and could lead to deleterious consequences such as increased ASD risk.

Our framework has several limitations. First, accurate estimation of direct and indirect effects requires all input GWAS to be sufficiently large with comparable sample sizes. Otherwise, the estimation performance will be limited by the least-powered GWAS. The current sample size in our EA analyses was not sufficient for significant association mapping or calculating well-powered PGS of direct and indirect EA effects. Second, we did not account for potential indirect sibling effects in our model. Although evidence has suggested that the indirect effects from siblings are negligible compared to direct effects and parental effects^15,16^, it may be important to account for sibling effects in certain applications^41^. Finally, our method provides unbiased estimates but will introduce a negative technical correlation between the direct and indirect effect estimates (**Supplementary Note**). This is not a unique issue in our approach and was also observed in analyses based on individual-level data^15,28^. Still, when sample size is limited, such technical correlations will shade the true genetic effects and hinder the interpretation of associations.

Taken together, our method has made important technical advancements in partitioning complex traits’ direct and indirect genetic effects. It provides statistically rigorous and computationally efficient estimates based on summary statistics from multi-generational GWAS, which provides a clear guidance on future study designs. If large genetic cohorts with multi-generational phenotypic information becomes the convention in the field, our method will have broad applications and can facilitate our understanding of the genetic basis of numerous human traits.

## Methods

### Method details

If genetic data are available in a number of parents-offspring trios, by regressing the offspring phenotype values *Y*_*M*_ on the offspring, maternal, and paternal genotypes (*G*_O,M,P_ respectively), the coefficients in joint regression estimate the direct genetic effect *β*_dir_, indirect maternal effect *β*_ind_mt_, and indirect paternal effect *β*_ind_pt_, respectively.

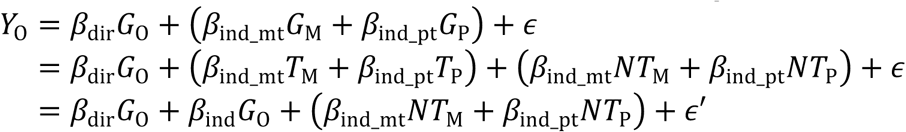

We define indirect genetic effect as the effect of a person’s genotype on his or her phenotype via the indirect pathway that goes through biological parents and the family environment. This effect is *β*_ind_mt_*T*_M_ + *β*_ind_pt_*T*_P_ in the equation above. Thus, the indirect effect size is obtained by regressing *β*_ind_mt_*T*_M_ + *β*_ind_pt_*T*_P_ on *G*_O_(= *T*_M_ + *T*_P_), which gives

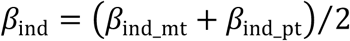

The estimated effect sizes from the marginal regression of *Y*_*M*_ on *G*_*M*_ (which is the standard GWAS) is given by 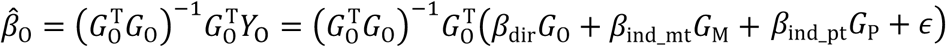. It is easy to find Cov(*G*_O_, *G*_M,P_) = *p*(1 - *p*)(1 + *α*), where *p* is the minor allele frequency (MAF) and *α* = Corr(*G*_M_, *G*_P_) is the correlation between spouses for the SNP. Then, we can express *β*_O_ in terms of the direct and indirect effect sizes. Similar expressions can also be derived for *β*_M_ and *β*_P_. Then using three equations, we can solve for the direct and indirect genetic effects analytically:

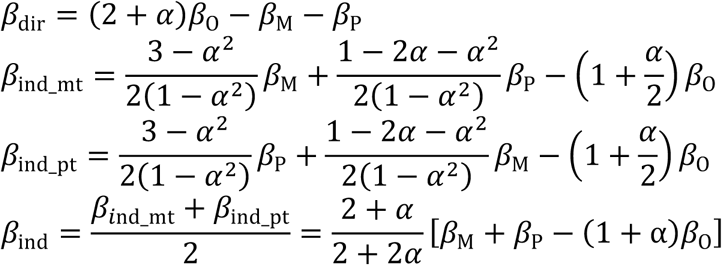

Our framework could also be naturally extended to the PGS level (**Supplementary Note**).

One technical note is we can see from the equation above that the covariance between 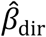 and 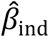 are always negative. This negative covariance should be reduced in magnitude as we increase the sample size since the variances of the input GWAS effect size estimates are reduced. Their correlation, however, will not reduce as we increase the sample size (**Supplementary Note**). This is because both the covariance and variance will decrease at a same rate as sample size increases. As a result, the correlation will remain at a fixed number if we keep the same ratios of the sample sizes.

### Simulation

We randomly selected 1,000 independent SNPs in the UKB data as causal variants and sampled their true effect sizes from a normal distribution with mean 0 and variance of 0.01^2^. We first simulated 3 sets of trios (sets 1-3). Each set consisted of 30,000 trios. In each trio, parental genotypes were independently simulated under binomial distributions using each SNP’s MAF. The offspring genotypes were generated from parental data following Mendelian inheritance. The offspring phenotype was computed as a weighted sum of these 1,000 SNPs plus a normal error term 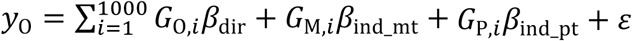. We used these 3 sets to run GWAS-O, GWAS-M, and GWAS-P, respectively. To simulate overlapping samples, we generated two additional sets (sets 4 and 5) of multi-generational families. In each family, we simulated data for 3 generations: 2 grandparents, 2 parents, and 1 child. Thus, these parents can be used as overlapping samples since we can compute both their own and their children’s phenotypes (**Supplementary Figures 1B** and **2**). *Var*(*ε*) was set to be 0.04 so that it accounted for ∼30% of the phenotypic variance.

We simulated 3 different scenarios: (1) GWAS-O, GWAS-M, and GWAS-P as inputs where all the samples in these 3 GWASs were independent; (2) GWAS-O and GWAS-MP as inputs where these 2 GWASs also used independent samples; (3) GWAS-O, GWAS-M, and GWAS-P as inputs, where GWAS-M and GWAS-P used independent samples, however all samples in GWAS-M and GWAS-P were also present in GWAS-O (**Supplementary Figure 2**). That is, for scenario (3), we used the parents in sets 4 and 5 to conduct GWAS-O, GWAS-M, and GWAS-P.

Among the 1,000 causal SNPs, we focused on one SNP with a MAF of 0.23. We used different settings for its true effect sizes: (*β*_dir_, *β*_ind_mt_, *β*_ind_pt_) = (0, 0, 0), (0, 0.02, 0.01), (0.02, 0, 0.01), (0.02, 0.02, 0), (0.02, 0.01, -0.01), (0.02, 0.01, 0.01), (0.02, 0.02, 0.01), and (0.02, -0.02, -0.01) to cover the null, positive direct and positive indirect, and positive direct and negative indirect effects combinations. For each setting, we repeated the simulation 1,000 times. For each repeat, we applied our method to estimate the direct and indirect effects. We also tested two other SNPs with MAF = 0.01 and 0.48 and the results looked similar.

### UK Biobank data processing

We used UKB data to conduct GWASs on birth weight and EA and perform PGS regression analyses. We excluded the participants that are recommended by UKB to be excluded from analysis (data field 22010 in the UKB), those with conflicting genetically inferred (data field 22001) and self-reported sex (data field 31), and those who withdrew from the study. UKB samples with European ancestry were identified from principal component analysis (data field 22006). PCs were computed using flashPCA2^42^. We used KING^43^ to infer the pairwise family kinship. We identified 154 pairs of monozygotic twins, 242 pairs of fraternal twins, 19,136 full sibling pairs, and 5,336 parent offspring pairs among 408,921 individuals with European ancestry in the UKB.

### Birth weight GWAS analysis

The own (N = 298,142) and maternal (N = 210,267) GWAS summary statistics reported in Warrington et al.^28^ were downloaded from the Early Growth Genetics Consortium website. We removed duplicated SNPs in each file, took SNP intersections between these two sets of summary statistics, and flipped the sign of effect size estimates when necessary such that the effective alleles were matched between the two input GWASs. We also downloaded the summary statistics for the inferred direct and indirect maternal genetic effects to compare with our results (**Supplementary Figure 6**). To have a fair comparison, we used the SNPs whose sample sizes and effective allele frequencies reported in the paper’s direct and indirect maternal effect summary statistics are consistent with those reported in their GWAS-O and GWAS-M summary statistics and heterogeneity *p*-value > 0.05. More than 8 million SNPs were used in the comparative analysis.

The UKB collected participants birth weight (data field 20022). Women who had at least one child were also asked for the birth weight of their first child (data field 2744). We also constructed two orthogonal phenotypes representing the direct and indirect components of birth weight following Warrington et al.^28^ The two new phenotypes were defined as: 2 (2*BW*_own_ – *BW*_offpring_)/3 and 2 (2*BW*_offspring_ - *BW*_own_)/3, where *BW*_own_ is own birth weight and *BW*_offspring_ is the offspring’s birth weight.

We conducted GWASs for these four phenotypes (i.e., own and offspring birth weights and the two orthogonal phenotypes) on 75,711 independent individuals of European ancestry who had both own and first child’s birth weights available. To compare, this number was 101,541 in Warrington et al.^28^ which included both Europeans and non-European samples. Birth weight was constructed following Warrington et al.^28^ Year of birth, genotype array, assessment center, and top 20 PCs were used as covariates. The results on own and first child’s birth weight were used as input in our framework to estimate the direct and maternal effects while the GWASs on orthogonal phenotypes were used as comparison.

### GWAS on offspring EA

We identified 5,336 parent-offspring pairs among the UKB EUR samples. Following Lee et al.^17^, we used the “qualification” (data field 6138) to compute the years of schooling as the EA phenotype. Year of birth, sex, genotype array, and top 20 PCs were used as covariates. We used parents from independent parent-offspring pairs with offspring EA phenotype and covariates information as GWAS samples. If both parents’ genotype data were available, we only included one of them in the analysis. The GWAS sample size N = 4,181 (2,619 females and 1,562 males).

In the HRS cohort, the respondent’s oldest child’s EA phenotype was constructed following Okbay et al.^44^ We kept only the independent (inferred by KING^43^) European parents (self-identified as “white/caucasian”) in our analyses. Year of birth, sex, and top 10 PCs were used as covariates. GWAS sample size N = 6,324 (3,780 females and 2,544 males).

In the WLS cohort, the oldest child’s education information was given by variables “z_rd01001”, “z_gd01001”, and “z_gd21001” corresponding to different rounds of collection. We used the maximum value whenever there was any inconsistency among different rounds. The EA phenotype was constructed following Lee et al.^17^ We required the GWAS samples to be of European ancestry (variable “z_ie008re”), independent (inferred by KING^43^), the oldest child was a biological offspring (“z_rd00401” and “z_gd00401”), the offspring’s EA was measured when the child was at least 30 years old and parent was at least 15 years older than the child. Year of birth, sex, and top 10 PCs were used as covariates. GWAS sample size N = 4,772 (2,513 females and 2,259 males).

PLINK^45^ version 1.9 was used to perform all GWASs. Finally, we meta-analyzed these three offspring EA GWASs using the inverse variance based approach in METAL^46^ to obtain the GWAS-MP as the input for our framework.

We also compared results given by our framework and SNIPar^15^ with a same set of data in UKB. Using the full siblings (N = 35,243 samples from 17,136 families) of European ancestries in UKB identified by KING^43^ (here, we only used the full siblings whose parents are not in the UKB), SNIPar imputed their expected average parental genotype. With the sum of imputed parental genotype and the observed offspring’s genotype jointly in the model, SNIPar computed the direct and indirect effects on EA with a linear mixed model (N = 34,956 samples from 17,135 unique families with non-missing EA phenotype). Using the same full sibling data, we performed GWAS-O using the observed siblings (N = 17,135 independent samples with phenotype and covariates available). Using the imputed sum of parental genotype, we ran GWAS-MP. Then our framework could also compute the direct and indirect effects using the two summary statistics and the comparison results are shown in **Supplementary Figure 14**. Year of birth, sex, genotyping array, and top 20 PCs were used as covariates in GWAS-O. In GWAS-MP, the offspring’s year of birth, genotyping array, and top 20 PCs were used as covariates where the PCs were computed using the imputed parental genotype by flashPCA2^42^.

### GWAS on own EA in UKB

We conducted GWAS-O (N = 356,719) using independent European samples in the UKB, excluding the full sibling samples (N = 35,243) that were used by SNIPar. The EA phenotypes were constructed following Lee et al^17^. Year of birth, sex, genotyping array, and top 20 PCs were used as covariates. We then meta-analyzed with EA3 summary statistics that excluded UKB samples. The reason for excluding the full siblings was because later we would do meta-analysis with SNIPar results which used the full sibling data.

### Genetic correlation analysis

We used both LDSC^29^ and GNOVA^31^ to compute genetic covariances and genetic correlations for any given pair of traits using their GWAS summary statistics. The results based on two approaches were comparable. The genetic correlation results shown in the main text were from LDSC. Details of the 45 traits used in the analysis and LDSC and GNOVA results are shown in **Supplementary Tables 5-7**.

### Polygenic score calculation and regression analysis

We performed PGS analysis on two sets of UKB samples with European ancestry: the first set was 16,580 pairs of full siblings and the second was 370,308 independent individuals. For each sample, two EA PGSs based on direct and indirect effect estimates were computed. To maximize the power and avoid overfitting, we used different input summary statistics to compute the direct and indirect effects. For the full sibling pairs, we first excluded full sibling pairs from the UKB samples, then used KING^43^ to identify a subset of independent individuals (N = 356,719) and ran an EA GWAS following Lee et al.^17^ We used METAL^46^ to meta-analyze it with EA3 GWAS that excluded 23andMe and UKB samples (N = 324,162). Together with the offspring EA GWAS as inputs, we computed the direct and indirect effect summary statistics which were used to compute the PGSs for the full sibling pairs in UKB. For the second set, we used the EA3 GWAS that excluded 23andMe and UKB samples and the offspring EA GWAS as input to estimate the direct and indirect effects.

To compute PGS, we first clumped the summary statistics in PLINK^45^ version 1.9 using the CEU samples in 1000 Genome Project Phase III cohort^47^ as the LD reference panel. We applied an LD window size of 1Mb and a pairwise r2 threshold of 0.1. Then, we computed PGS using PRSice-2^48^ with a fine-tuned p-value cutoff given by PUMAS^32^. PUMAS uses GWAS summary statistics as input and output an optimal p-value cutoff that gives the highest *R*^2^ for the PGS regression analysis. Since the PGS will use only the SNPs that are present in the target samples, we used only the SNPs that are present in the summary statistics, LD reference panel, and the target samples when running PUMAS.

We used software R^49^ version 3.5.1 to run linear regression of EA on PGSs. Both the EA phenotype and PGS were standardized. For full sibling pairs, we regressed EA difference between siblings on PGS differences. For independent samples, we used year of birth, sex, genotype array, assessment center, and top 10 PCs as covariates. *R*^2^ was computed as the ratio of sum squares by PGS to the total sum of squares.

### pTDT analysis

Three ASD cohorts were used in the pTDT analysis: AGP (N = 2,188 trios), SSC (1,794 proband trios and 1,430 sibling trios), and SPARK (3,822 proband trios and 1,812 sibling trios). Details of data processing in these cohorts have been described previously^38^. To compute PGS, we first used PLINK^45^ version 1.9 to clump the direct and indirect effect summary statistics using the CEU samples in 1000 Genome Project Phase III cohort^47^ as the LD reference panel. We applied an LD window size of 1Mb and a pairwise r2 threshold of 0.1. PGSs were computed using PRSice-2^48^ with optimal p-value cutoffs estimated by PUMAS^32^. We performed pTDT^35^ to measure the transmission disequilibrium in EA polygenic risks for ASD probands and siblings.

### URL

AGP (https://www.ncbi.nlm.nih.gov/projects/gap/cgi-bin/study.cgi?study_id=phs000267.v5.p2);

SSC (https://www.sfari.org/resource/simons-simplex-collection/);

SPARK (https://www.sfari.org/resource/spark/);

Early Growth Genetics Consortium (https://egg-consortium.org/birth-weight-2019.html);

SNIPar (https://github.com/AlexTISYoung/SNIPar)

## Data and code availability

The DONUTS package is available at https://github.com/qlu-lab/DONUTS

## Acknowledgements

This project was supported by the Clinical and Translational Science Award (CTSA) program, through the NIH National Center for Advancing Translational Sciences (NCATS), grant UL1TR000427. We also acknowledge research support from the University of Wisconsin-Madison Office of the Chancellor and the Vice Chancellor for Research and Graduate Education with funding from the Wisconsin Alumni Research Foundation and the Waisman Center pilot grant program at the University of Wisconsin-Madison. We are grateful to all the families participating in the Autism Genome Project (AGP), the Simons Simplex Collection (SSC), and the Simons Foundation Powering Autism Research for Knowledge (SPARK) study. We thank Dr. Aysu Okbay for providing the EA meta-analysis results with UKB data removed. We thank Drs. Jan Greenberg and Marsha Mailick for their assistance in WLS data collection and processing.

## Author contribution

Q.L. conceived and designed the study.

Y.W. and Q.L. developed the statistical framework.

Y.W. performed simulations and data analysis.

X.Z. assisted the data analysis.

Y.L. assisted developing the statistical framework and simulation.

Z.Z. calculated and fine-tuned polygenic scores.

J.C. performed pTDT analysis.

B.Z and J.F. assisted HRS data preparation and interpretation.

J.L. assisted WLS data collection and processing and advised on neurodevelopmental disorder genetics.

Q.L. advised on statistical and genetic issues.

Y.W. and Q.L. wrote the manuscript.

All authors contributed in manuscript editing and approved the manuscript.

