## Supplementary Figure for "Estimating genetic nurture with summary statistics of multi-generational genome-wide association studies"

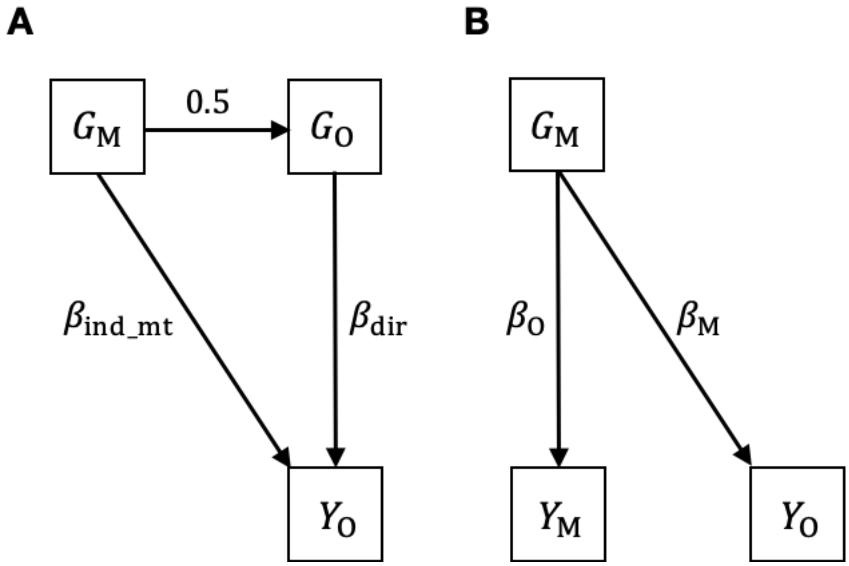


**Supplementary Figure 1**. **Some special cases in our statistical framework. (A)** Our full model presented in **Figure 1** could be simplified if the indirect genetic effect from one parent is 0. **(B)** demonstrates the sample overlapping case: a mother reported both her own phenotype $Y_{\mathrm{M}}$ and her offspring's phenotype $Y_{\mathrm{O}}$, thus she will be present in both GWAS-O and GWAS-M, leading to correlated $\hat{\beta}_{\mathrm{O}}$ and $\hat{\beta}_{\mathrm{M}}$.

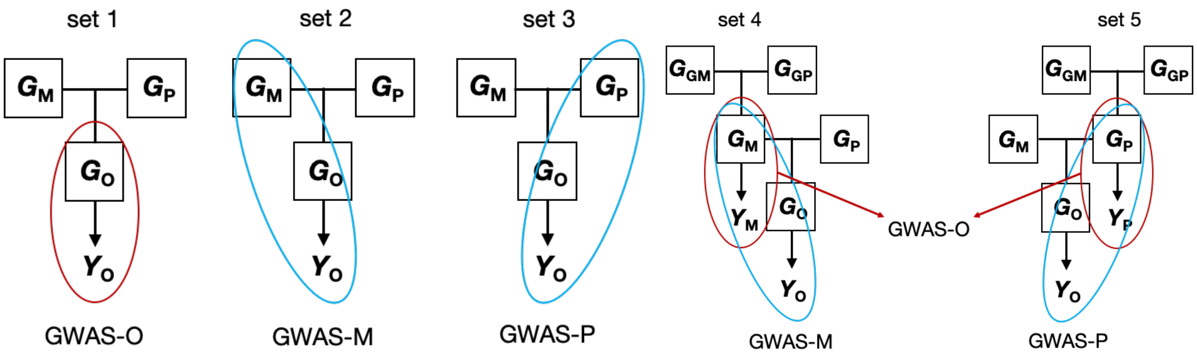


**Supplementary Figure 2. Simulation designs for GWAS-O, GWAS-M, and GWAS-P**. Simulated sets 1-3 consisted of trios whereas each simulated family had 3 generations in sets 4 and 5. Sets 1-3 are used to conduct GWAS-O, GWAS-M, and GWAS-P. For GWAS-MP, we pooled mothers in set 2 and fathers in set 3 to run the GWAS. Parents in sets 4 and 5 are used as overlapping samples since they have both their own and their children's phenotype.

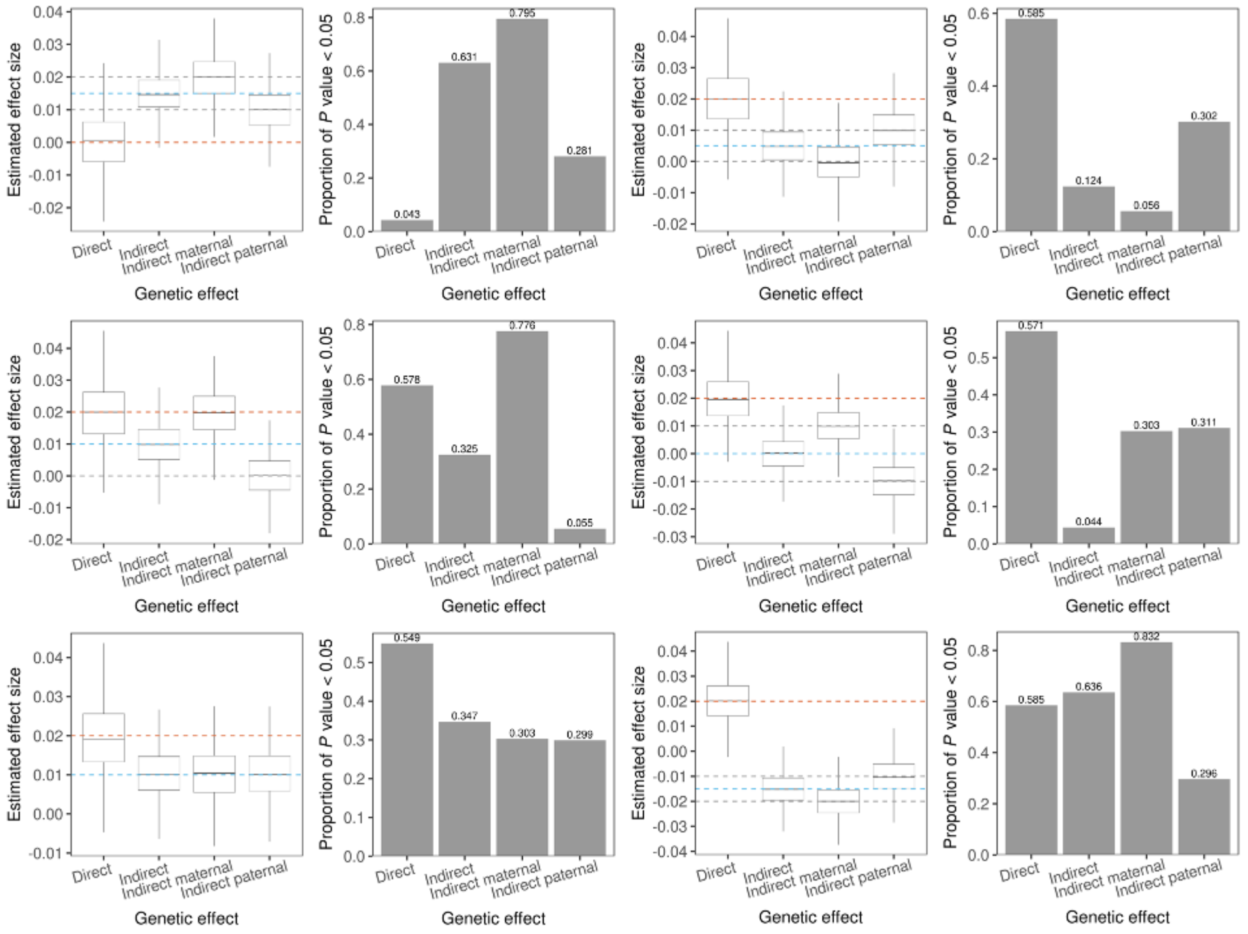


**Supplementary Figure 3. Additional simulation results when summary statistics from independent GWAS-O, GWAS-M, and GWAS-P are used as input.** $n_{\mathrm{O}} = n_{\mathrm{M}} = n_{\mathrm{P}} = 30\mathrm{K}$ in each simulation. Red and blue dashed lines indicate the true direct and indirect effect size, respectively. The grey dashed lines indicate the true indirect maternal and paternal effect sizes. True effect sizes $(\beta_{\mathrm{dir}}, \beta_{\mathrm{ind\_mt}}, \beta_{\mathrm{ind\_pt}})$ = (0, 0.02, 0.01), (0.02, 0, 0.01), (0.02, 0.02, 0), (0.02, 0.01, -0.01), (0.02, 0.01, 0.01), and (0.02, -0.02, -0.01) for the above 6 sets of figures (top to bottom, left to right), respectively. Box plots show the effect size estimates for the 1,000 SNPs used in simulation and the bar plots show the proportion of *p* values smaller than 0.05.

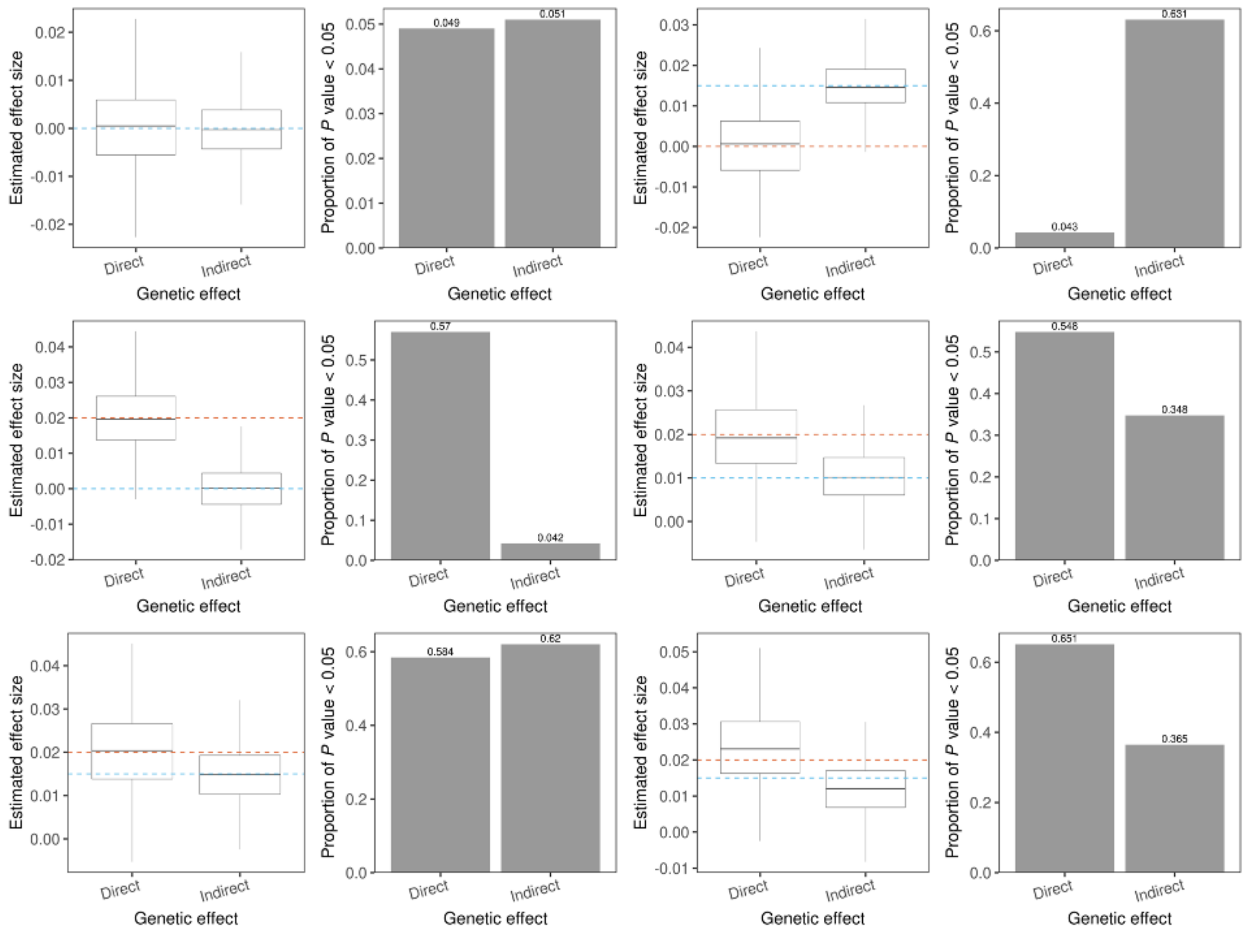


**Supplementary Figure 4. Additional simulation results when summary statistics from independent GWAS-O and GWAS-MP are used as input.** Legends have the same meanings as those in **Supplementary Figure 3**. Effect sizes $(\beta_{\mathrm{dir}}, \beta_{\mathrm{ind\_mt}}, \beta_{\mathrm{ind\_pt}})$ = (0, 0, 0), (0, 0.02, 0.01), (0.02, 0.01, -0.01), (0.02, 0.01, 0.01), (0.02, 0.02, 0.01), and (0.02, 0.02, 0.01) for the above 6 sets of plots (top to bottom, left to right), respectively. Sample sizes for GWAS-M and GWAS-P are $n_{\mathrm{M}} = n_{\mathrm{P}} = 30\mathrm{K}$ for the first 5 plots and $n_{\mathrm{O}} = n_{\mathrm{P}} = 30\mathrm{K}, n_{\mathrm{M}} = 15\mathrm{K}$ for the last plot. Therefore, for the last plot we have $\beta_{\mathrm{ind\_mt}} \neq \beta_{\mathrm{ind\_pt}}$ and $n_{\mathrm{M}} \neq n_{\mathrm{P}}$. As a result, the effect size estimates are biased as shown in the last plot.

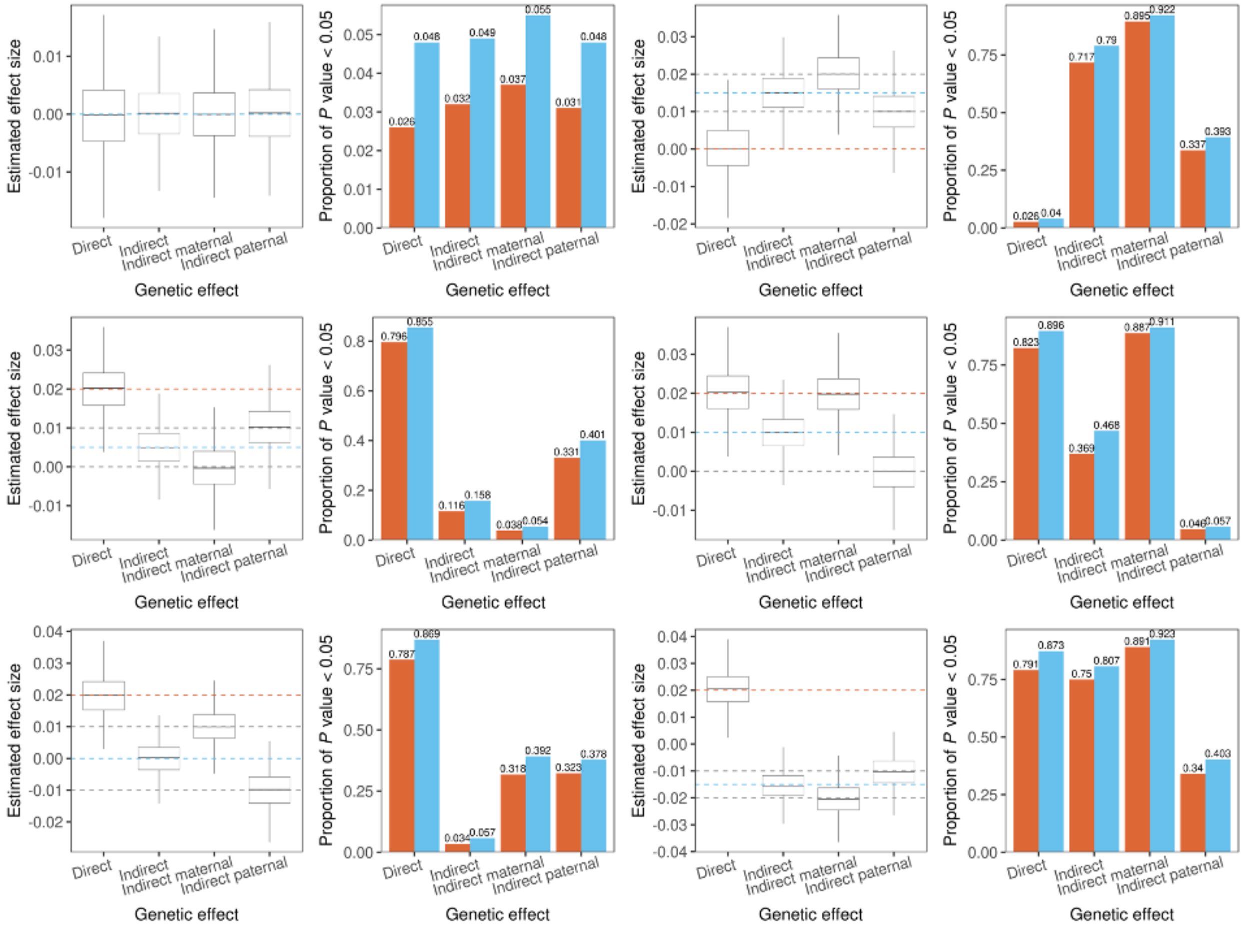


**Supplementary Figure 5. Additional simulation results when input GWAS have complete sample overlaps.** GWAS-O, GWAS-M, and GWAS-P were used as input where all the samples in GWAS-M and GWAS-P were also used in GWAS-O (**Supplementary Figure 2**). Blue and red bars show the statistical power with and without sample overlap correction, respectively. Other legends have the same meanings as those in **Supplementary Figure 3**. Effect sizes $(\beta_{\text{dir}}, \beta_{\text{ind\_mt}}, \beta_{\text{ind\_pt}})$ = (0, 0, 0), (0, 0.02, 0.01), (0.02, 0, 0.01), (0.02, 0.02, 0), (0.02, 0.01, -0.01), and (0.02, -0.02, -0.01) for the above 6 sets of figures (top to bottom, left to right), respectively. $n_{\text{M}} = n_{\text{P}} = 30\text{K}, n_{\text{O}} = n_{\text{M}} + n_{\text{P}} = 60\text{K}$.

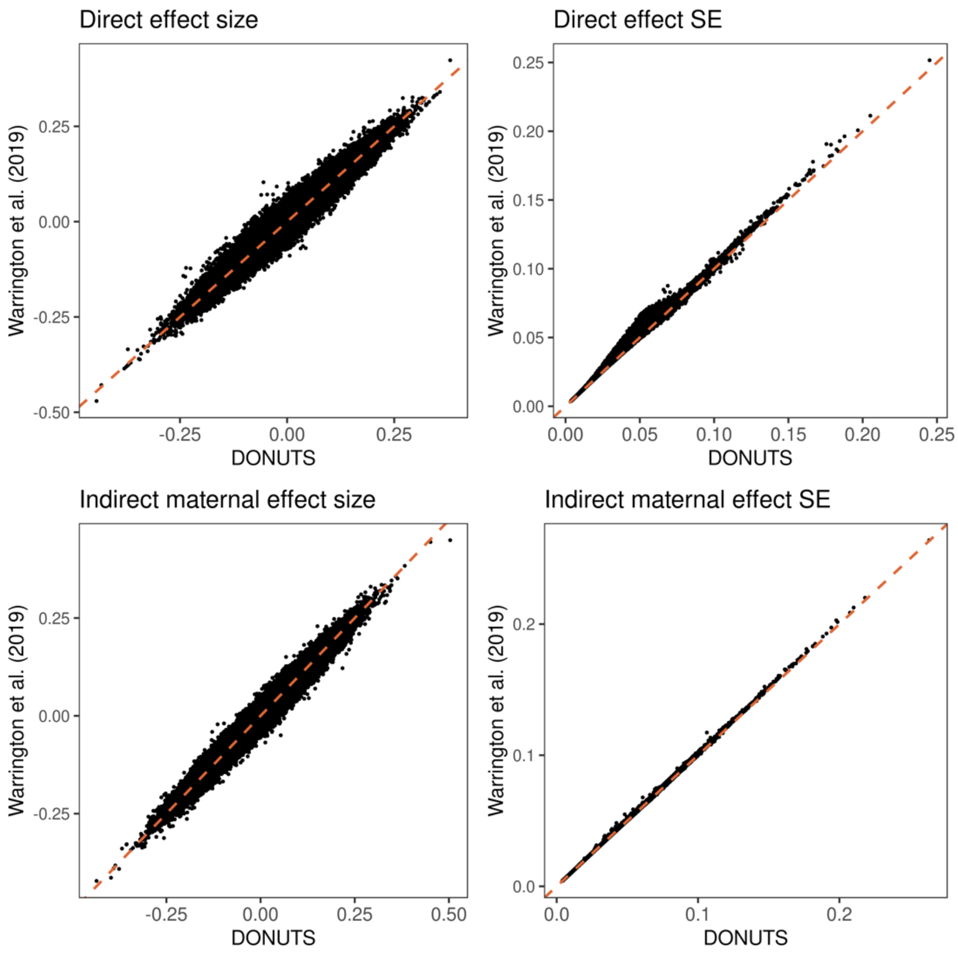


**Supplementary Figure 6. Comparisons of effect sizes and standard errors given by DONUTS with those obtained from Warrington *et al*.** Effects sizes and standard errors are highly consistent between two analyses, although standard errors from Warrington *et al*. showed a minor inflation due to sample overlaps that were not fully accounted for. Effect size estimates are not affected by overlapping samples.

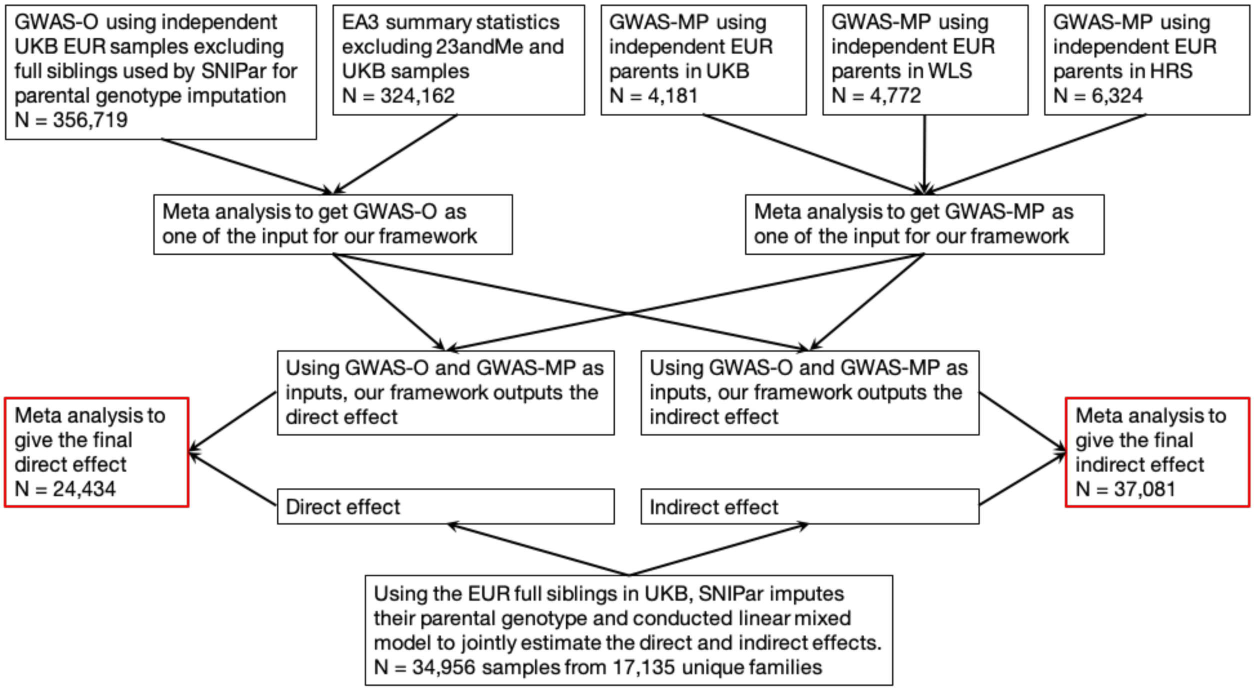


**Supplementary Figure 7. Flowchart of the analysis to obtain the direct and indirect genetic effects on EA.** Final results used in the main text are outlined by red boxes. An alternative approach to compute the direct/indirect effects is to use EA3_excl23andMe and meta-analyzed GWAS-MP based on UKB, WLS, HRS and SNIParFS, where for SNIParFS we used SNIPar to impute UKB full sibling's parental genotypes and then run GWAS-MP. Note, in this approach there are overlapping samples between GWAS-O and GWAS-MP, however, DONUTS could account for it. This approach is less powerful (effective sample sizes are 25,925 and 26,586 for the direct and indirect effects, respectively; **Supplementary Table 3**) and the results are also consistent with the results used in the main text: LDSC genetic correlations between these two approaches are 3.63 and 1.28 for the direct and indirect effects, respectively. (GNOVA genetic correlations are 2.50 and 0.89 for the direct and indirect effects, respectively.)

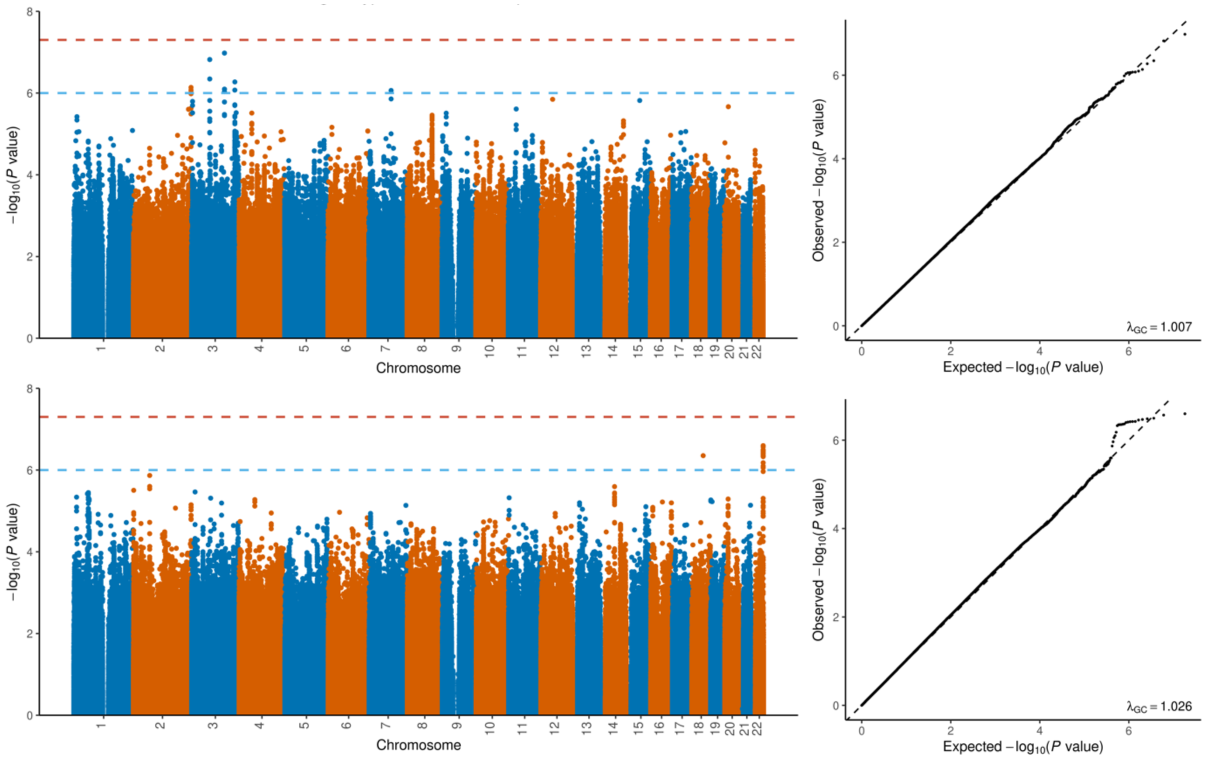


**Supplementary Figure 8. Manhattan (1st column) and Quantile-Quantile (2nd column) plots for direct genetic effect (1st row) and indirect genetic effect (2nd row) on EA.**

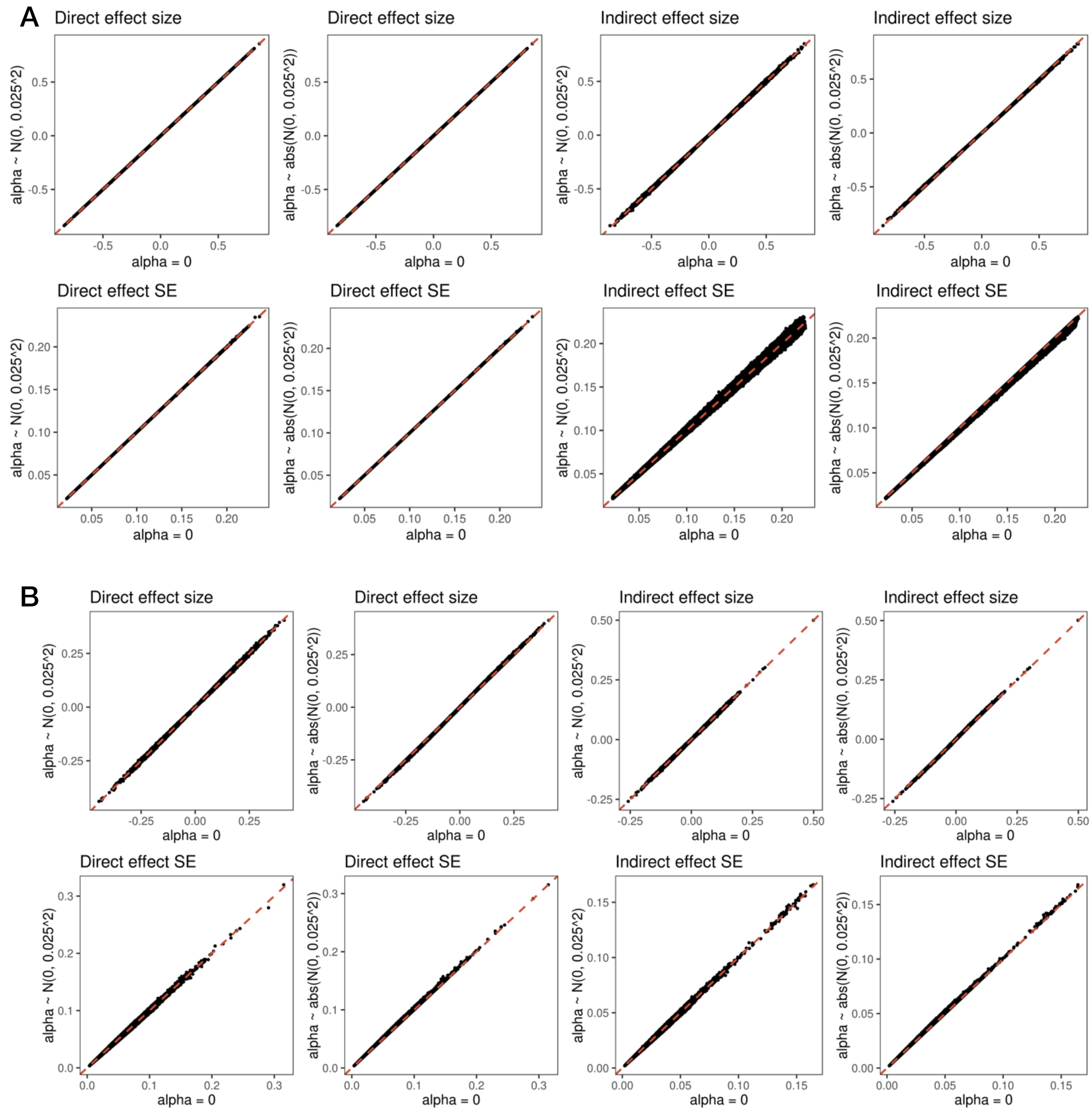


**Supplementary Figure 9. Impact of assortative mating on the direct and indirect effect estimates.** (**A**) EA; (**B**) birth weight. We simulated $\alpha \sim N(0, 0.025^2)$ to have comparable values with those from real couples. We also compared the case when all $\alpha$'s are positive. X-axis shows results when assuming $\alpha = 0$; Y-axis shows results based on non-zero $\alpha$ values. Each data point is a SNP. To obtain the direct and indirect effects in **A**, we used EA GWAS summary statistics from Lee *et al*. as GWAS-O (N = 766,345; 23&me samples excluded) and the meta analyzed GWAS-MP (N = 15,277) based on UKB, WLS, and HRS samples.

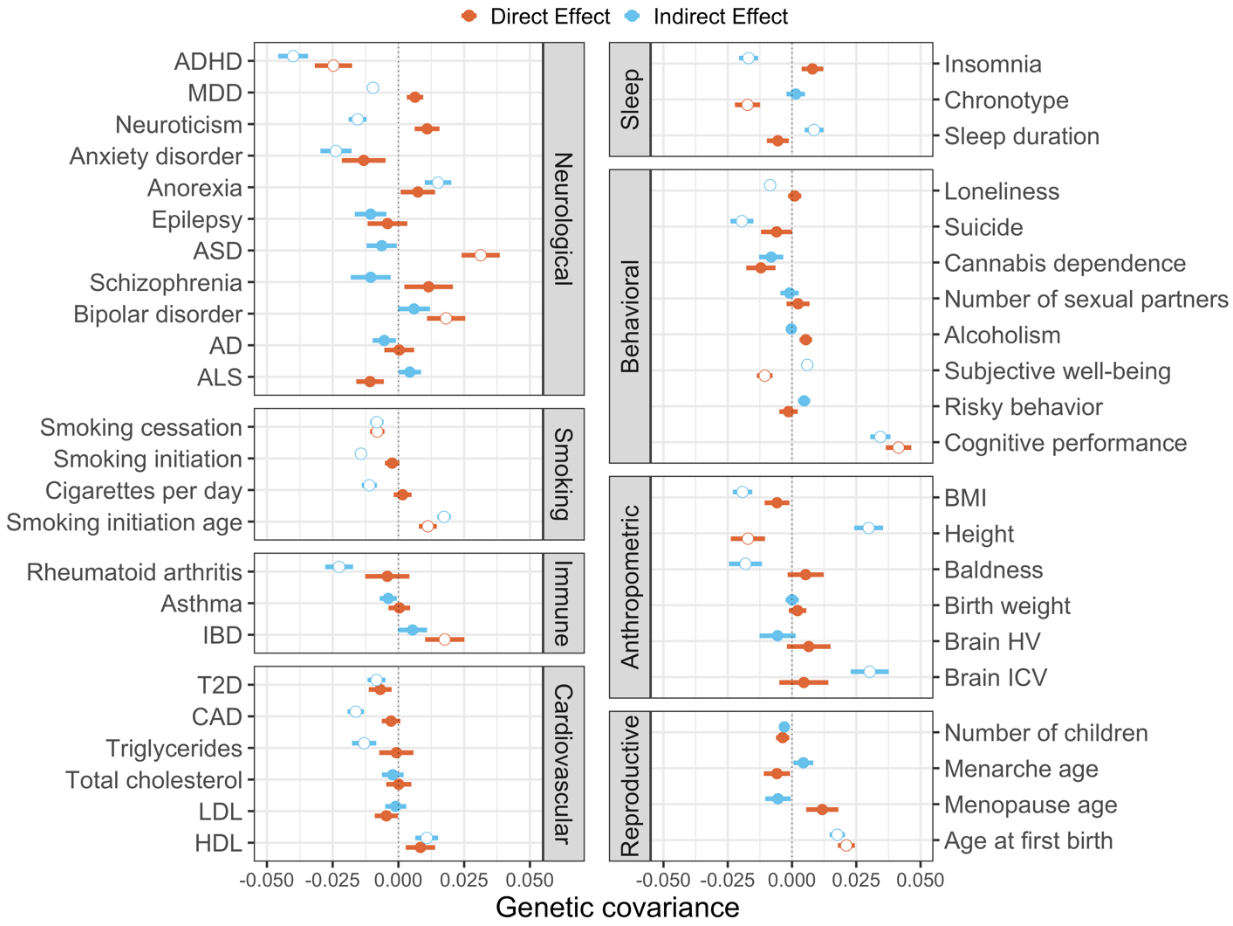


**Supplementary Figure 10. Genetic covariances of EA (direct and indirect effects) with 45 complex traits estimated by GNOVA.** Dots and intervals indicate the point estimates and standard error of genetic covariances, respectively. Significant correlations at an FDR cutoff of 0.05 are highlighted with white circles. ADHD: attention deficit/hyperactivity disorder; MDD: major depressive disorder; ASD: autism spectrum disorder; AD: Alzheimer's diseases; ALS: amyotrophic lateral sclerosis; IBD: inflammatory bowel disease; T2D: type-2 diabetes; CAD: coronary artery disease; LDL and HDL: low and high-density lipoprotein; BMI: body-mass index; HV: hippocampal volume; ICV: intracranial volume.

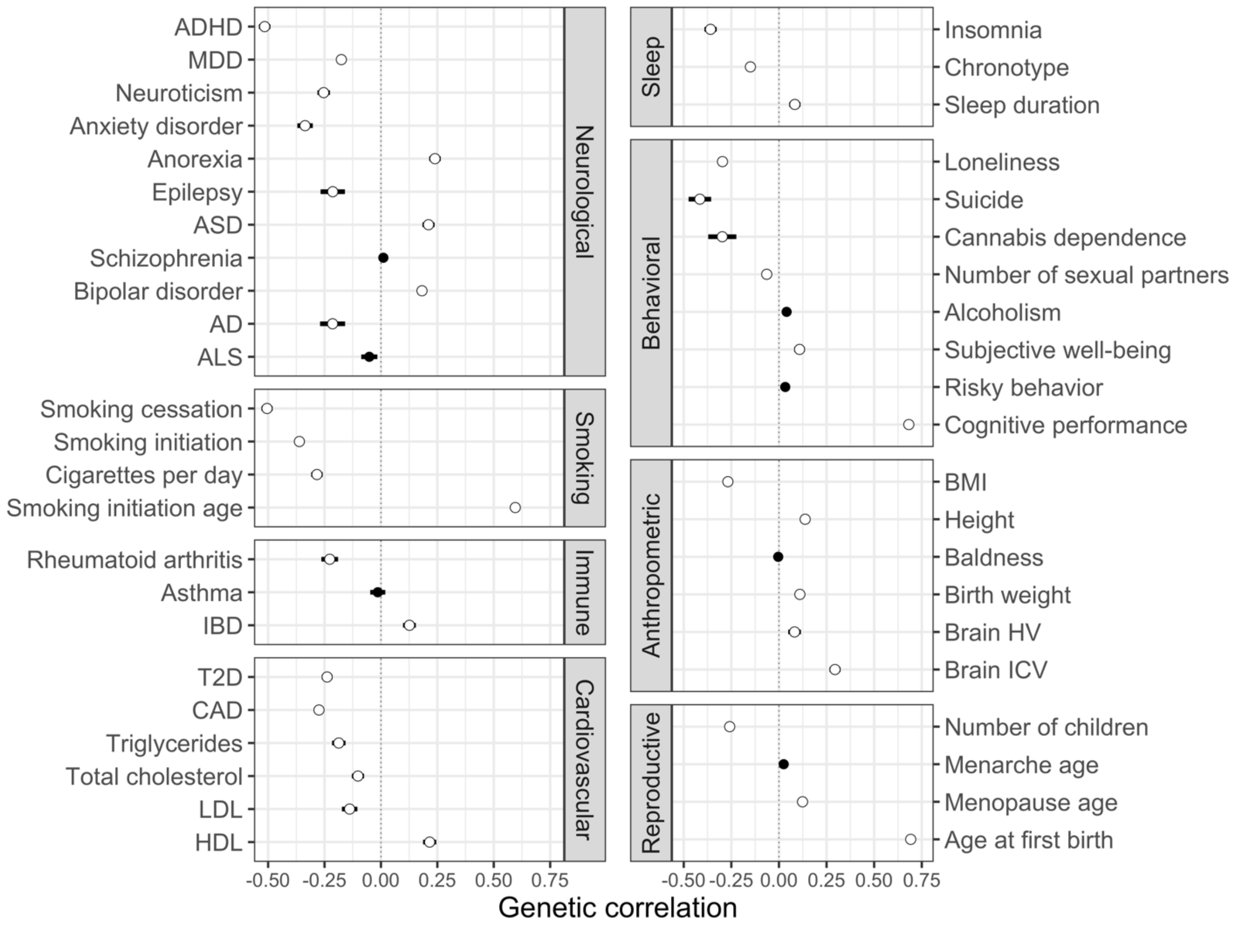


**Supplementary Figure 11. Genetic correlations of EA with 45 complex traits estimated by LDSC.** Dots and intervals indicate the point estimates and standard error of genetic correlations, respectively. Significant correlations at an FDR cutoff of 0.05 are highlighted with white circles. Out of 45 traits, 38 are significantly correlated with EA. ADHD: attention deficit/hyperactivity disorder; MDD: major depressive disorder; ASD: autism spectrum disorder; AD: Alzheimer's diseases; ALS: amyotrophic lateral sclerosis; IBD: inflammatory bowel disease; T2D: type-2 diabetes; CAD: coronary artery disease; LDL and HDL: low and high-density lipoprotein; BMI: body-mass index; HV: hippocampal volume; ICV: intracranial volume.

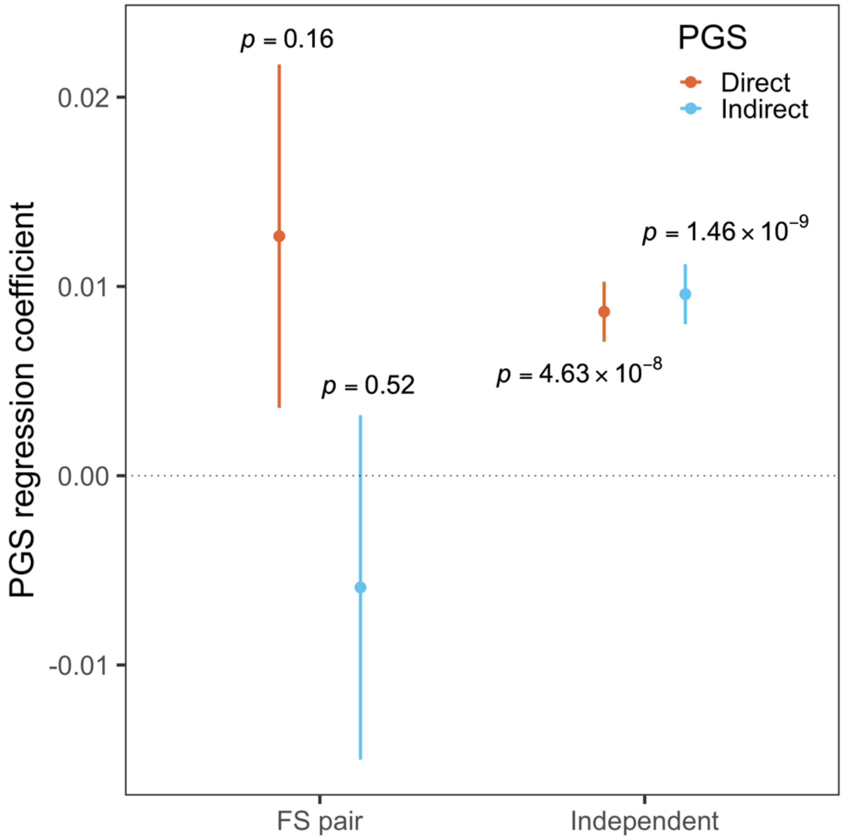


**Supplementary Figure 12. Predictive performances of direct and indirect PGS for EA evaluated using UKB participants with European (EUR) ancestry.** We generated bioinformatically fine-tuned PGS for direct and indirect components of EA. The PGS effect size was obtained by regressing EA on direct and indirect PGS. Dots and bars represent the point estimates and standard error, respectively. There were 16,580 full sibling (FS) pairs and 370,308 independent individuals used in the regression analysis. Results are also included in **Supplementary Table 8.**

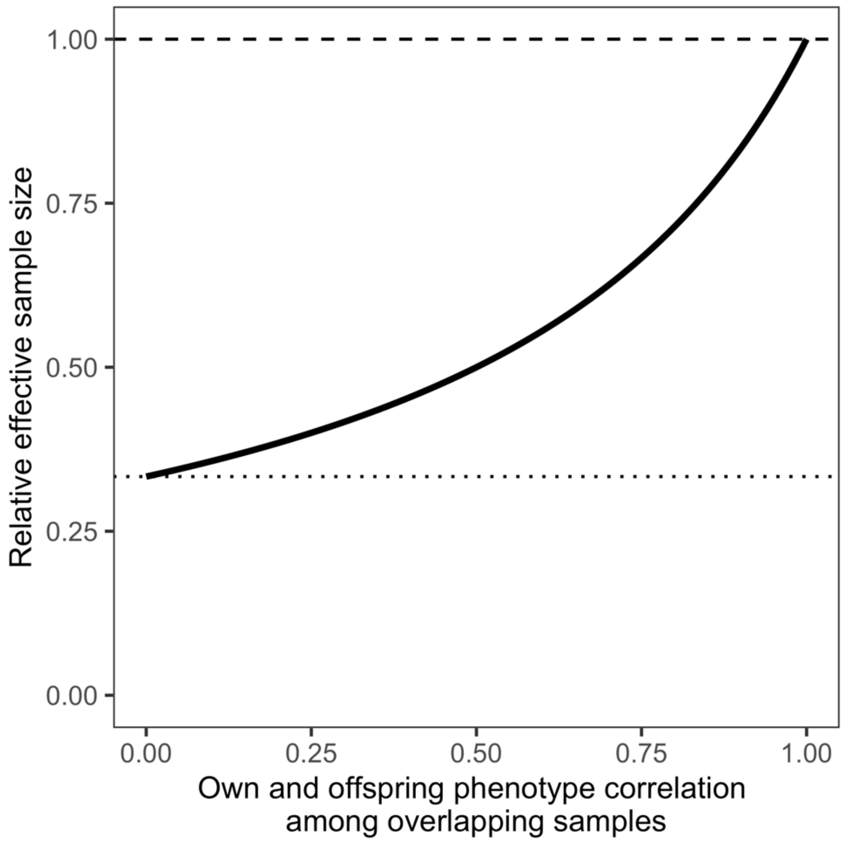


**Supplementary Figure 13. Relative effective sample sizes between our design and a trio-based design for direct and indirect effects as a function of phenotype correlation among overlapping samples in the input GWAS.** We could get the empirical effective sample sizes for the direct and indirect genetic effects from either a trio-based design with individual-level genetic data or using DONUTS on multi-generational GWAS (**Supplementary Note**). We compared their effective sample sizes and the y-axis shows the ratio of the effective sample size from our study design to that from a trio design. The x-axis is the phenotypic correlation among overlapping samples. Our effective sample sizes are functions of sample sizes from each input GWAS, number of overlapping samples, and phenotype correlation among overlapping samples. In the comparison, we fixed the number of total individuals to be $3N$ for each design: $N$ trios for the trio design vs. $n_O = n_M = n_P = N$ for GWAS-O,M,P, respectively in our approach. Half of GWAS-O samples were also in GWAS-M while the other half were in GWAS-P. For simplicity, we assumed $\alpha = 0$ in this analysis. The results were the same for either the direct or indirect effects. When there is no sample overlap or if the phenotype correlation is 0, our effective sample size is 3 times smaller (dotted horizontal line) than that from a trio design. When the phenotype correlation approaches 1, our approach could be as powerful as the trio design. When using GWAS-O and GWAS-MP as input (i.e., case ii in **Table 1**), if $n_O = N, n_{MP} = 2N$ and all the GWAS-O samples are also in GWAS-MP (i.e., every individual reports both self and offspring's phenotype), we'll get the same results as shown in this figure.

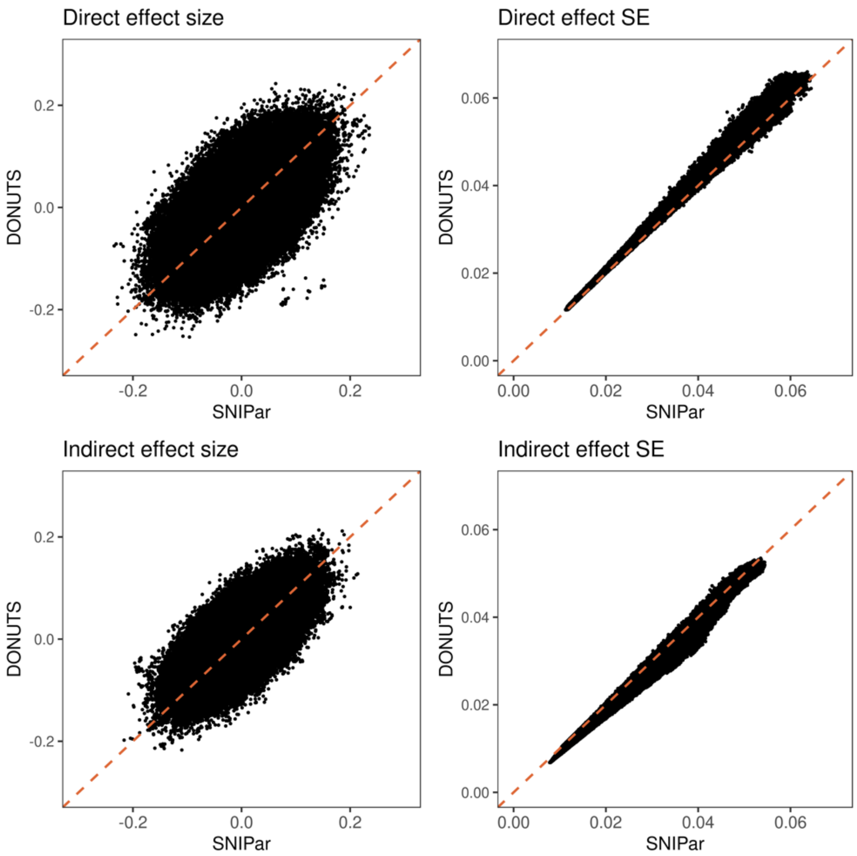


**Supplementary Figure 14. Direct (1st row) and indirect effects (2nd row) given by DONUTS (y-axis) vs. SNIPar (x-axis) using a same set of data.** Using the full siblings (N = 35,243 samples from 17,136 families) in UKB, SNIPar imputed their expected average parental genotype. With the imputed sum of parental genotype and the observed offspring's genotype jointly in the model, SNIPar computed the direct and indirect effects on EA using a linear mixed model (N = 34,956 samples from 17,135 unique families with non-missing EA phenotype). Using the same data, we performed GWAS-O using the observed siblings (N = 17,135 independent samples with phenotype and covariates available). Using the sum of the imputed parental genotype, we performed GWAS-MP. Then, our framework could also compute the direct and indirect effects using the two summary statistics. Additional details of this analysis are described in **Methods** and **Supplementary Note.**
