## Supplementary Note for "Estimating genetic nurture with summary statistics of multi-generational genome-wide association studies"

### Covariances among genotypes within families

If assortative mating exists, parental genotypes will be correlated. Define the correlation between spouses at a locus

$$\mathrm{Corr}(G_M, G_P) = \frac{\mathrm{Cov}(G_M, G_P)}{2p(1-p)} = \alpha,$$

where $G_{M,P}$ are the maternal and paternal genotypes, respectively. $p$ is the minor allele frequency (MAF) of the single-nucleotide polymorphisms (SNP). $\alpha$ quantifies the assortative mating at SNP level. The variances of the parental genotype are

$$\mathrm{Var}\left(G_{M,P}\right) = 2p(1-p)$$

Since half of the parental genotypes are randomly transmitted to their offspring, we can re-write $G_{M,P} = T_{M,P} + NT_{M,P}$, where *T* and *NT* represent the transmitted and non-transmitted alleles. Their offspring's genotype is $G_O = T_M + T_P$, the variances and covariances within a family can be derived:

$$\mathrm{Cov}(G_M, G_P) = \mathrm{Cov}(T_M + NT_M, T_P + NT_P) = 4\mathrm{Cov}(T_M, T_P) = 2p(1-p)\alpha$$
$$\mathrm{Cov}(T_M, T_P) = p(1-p)\alpha/2$$
$$\mathrm{Var}(G_O) = \mathrm{Var}(T_M + T_P) = \mathrm{Var}(T_M) + \mathrm{Var}(T_P) + 2\mathrm{Cov}(T_M, T_P) = 2p(1-p)\left(1 + \frac{\alpha}{2}\right)$$
$$\begin{aligned} Cov(G_O, G_M) &= Cov(T_M + T_P, T_M + NT_M) = Var(T_M) + Cov(T_P, T_M) + Cov(T_P, NT_M) \\ &= p(1-p)(1+\alpha) \\ &= Cov(G_O, G_P) \end{aligned}$$

### SNP level direct and indirect effects dissection

#### Full model using trio genetic data

If genetic data are available in a number of parents-offspring trios, by regressing the offspring phenotype values $Y_O$ on the offspring, maternal, and paternal genotypes, the coefficients in joint regression estimate the direct genetic effect $\beta_{\text{dir}}$, indirect maternal effect $\beta_{\text{ind\_mt}}$, and indirect paternal effect $\beta_{\text{ind\_pt}}$, respectively.

$$Y_{\text{O}} = \beta_{\text{dir}} G_{\text{O}} + \beta_{\text{ind\_mt}} G_{\text{M}} + \beta_{\text{ind\_pt}} G_{\text{P}} + \epsilon$$

To be consistent with the definition of the direct genetic effect, we define indirect genetic effect as the effect of a person's genotype $G_0$ on her phenotype via the indirect pathway that goes through biological parents and the family environment. This effect is $\beta_{\text{ind\_mt}} T_{\text{M}} + \beta_{\text{ind\_pt}} T_{\text{P}}$ in the above equation since it is due to $G_O = T_M + T_P$. The indirect effect size is obtained by regressing it on $G_{\text{O}}$ which gives

$$\begin{aligned}\hat{\beta}_{ind} &= \frac{G_O^T\left(\beta_{ind\_mt} T_M + \beta_{ind\_pt} T_P\right)}{G_O^T G_O} \\ &\rightarrow \left(\beta_{ind\_mt} + \beta_{ind\_pt}\right) \frac{p(1-p)(1+\alpha/2)}{2p(1-p)(1+\alpha/2)} \\ &= \frac{\beta_{ind\_mt} + \beta_{ind\_pt}}{2}\end{aligned}$$

Thus, we have

$$\beta_{ind} = \left(\beta_{ind\_mt} + \beta_{ind\_pt}\right)/2$$

### Another definition of $\boldsymbol{\beta_{ind}}$

Although the above seems to be a natural and coherent way to define the indirect effect, another possible definition is to use the total nurture effect $\beta_{\text{ind\_mt}} G_{\text{M}} + \beta_{\text{ind\_pt}} G_{\text{P}}$. Let's use $\beta_{\text{ind}}^*$ to denote it. Then,

$$\begin{aligned}\hat{\beta}_{ind}^* &= \frac{G_O^T\left(\beta_{ind\_mt} G_M + \beta_{ind\_pt} G_P\right)}{G_O^T G_O} \\ &\rightarrow \left(\beta_{ind\_mt} + \beta_{ind\_pt}\right) \frac{p(1-p)(1+\alpha)}{2p(1-p)(1+\alpha/2)} \\ &= \frac{\beta_{ind\_mt} + \beta_{ind\_pt}}{2} \frac{1+\alpha}{1+\alpha/2}\end{aligned}$$

Under this definition,

$$\beta_{ind}^{*} = \frac{\beta_{ind\_mt} + \beta_{ind\_pt}}{2} \frac{1+\alpha}{1+\alpha/2} = \beta_{ind} \frac{1+\alpha}{1+\alpha/2}$$

and when $\alpha = 0$, $\beta_{ind}^{*} = \beta_{ind}$. The reason for the difference between these two definitions is due to assortative mating such that $T_M$ and $NT_P$, $T_P$ and $NT_M$ become correlated. Throughout our entire analysis, we used our first definition for the indirect effects of a SNP since it (i.e., $\beta_{ind} = (\beta_{ind\_mt} + \beta_{ind\_pt})/2$) also naturally appears in many derivations.

**Direct and indirect effects estimated from marginal regressions**

Consider the following three marginal GWAS of three independent samples:

GWAS-O: regression of own phenotype on own genotype $Y_{\mathrm{O}} = G_{\mathrm{O}}\beta_{\mathrm{O}} + u_{\mathrm{O}}$

GWAS-M: regression of offspring phenotype on maternal genotype $Y_{\mathrm{O}} = G_{\mathrm{M}}\beta_{\mathrm{M}} + u_{\mathrm{M}}$

GWAS-P: regression of offspring phenotype on paternal genotype $Y_{\mathrm{O}} = G_{\mathrm{P}}\beta_{\mathrm{P}} + u_{\mathrm{P}}$

The estimated effect sizes are

$$\begin{aligned}\hat{\beta}_O &= \frac{G_O^T Y_O}{G_O^T G_O} = \frac{G_O^T(\beta_{dir}G_O + \beta_{ind\_mt}G_M + \beta_{ind\_pt}G_P + \epsilon_O)}{G_O^T G_O} \\ &= \beta_{dir} + \beta_{ind\_mt}\frac{G_O^T G_M}{G_O^T G_O} + \beta_{ind\_pt}\frac{G_O^T G_P}{G_O^T G_O} + \frac{G_O^T \epsilon_O}{G_O^T G_O}\end{aligned}$$

By law of large numbers, as GWAS-O sample size $n_O \to \infty$

$$\frac{1}{n_O} G_O^T G_M = \frac{1}{n_O}\sum_{i=1}^{n_O} G_{O,i}\, G_{M,i} \xrightarrow{p} E(G_{O,i}G_{M,i}) = Cov(G_{O,i}, G_{M,i}) = p(1-p)(1+\alpha)$$

Similarly,

$$\frac{1}{n_O} G_O^T G_O \to 2p(1-p)(1+\alpha/2),$$

$$\frac{1}{n_O} G_O^T G_P \to p(1-p)(1+\alpha),$$

$$\frac{1}{n_O} G_O^T \epsilon_O \rightarrow 0.$$

Then by Slutsky's Theorem,

$$\frac{G_O^T G_M}{G_O^T G_O} \xrightarrow{d} \frac{p(1-p)(1+\alpha)}{2p(1-p)(1+\alpha/2)} = \frac{1+\alpha}{2+\alpha}$$

Thus,

$$\hat{\beta}_0 \xrightarrow{d} \beta_{dir} + \frac{1+\alpha}{2+\alpha}\left(\beta_{ind\_mt} + \beta_{ind\_pt}\right) = \beta_{dir} + \left(1 + \frac{\alpha}{2+\alpha}\right)\beta_{ind}$$

Since $G_O$ is centered, $G_O^T G_O$ has lower bound greater than $0$. When the phenotype is also bounded, $\hat{\beta}_O$ is bounded. Since $\hat{\beta}_O$ is unbiased estimator of $\beta_O$, by the Lebesgue's dominated convergence theorem,

$$\beta_O = \lim \beta_O = \lim E(\hat{\beta}_O) = E(\lim \hat{\beta}_O) = \beta_{dir} + \frac{1+\alpha}{2+\alpha}(\beta_{ind\_mt} + \beta_{ind\_pt})$$

Similarly, from the other two GWASs

$$\beta_M = \frac{1+\alpha}{2}\beta_{dir} + \beta_{ind\_mt} + \alpha\beta_{ind\_pt}$$

$$\beta_P = \frac{1+\alpha}{2}\beta_{dir} + \alpha\beta_{ind\_mt} + \beta_{ind\_pt}$$

It is clear that the effect size estimates from a marginal regression is a combination of both direct and indirect effects and is also affected by assortative mating.

Solving for $\beta_{dir}, \beta_{ind\_mt}, \beta_{ind\_pt}$:

$$\beta_{dir} = (2+\alpha)\beta_O - \beta_M - \beta_P$$

$$\beta_{ind\_mt} = -\left(1 + \frac{\alpha}{2}\right)\beta_O + \frac{3-\alpha^2}{2(1-\alpha^2)}\beta_M + \frac{1-2\alpha-\alpha^2}{2(1-\alpha^2)}\beta_P$$

$$\beta_{ind\_pt} = -\left(1 + \frac{\alpha}{2}\right)\beta_O + \frac{1-2\alpha-\alpha^2}{2(1-\alpha^2)}\beta_M + \frac{3-\alpha^2}{2(1-\alpha^2)}\beta_P$$

Then, the indirect effect

$$\beta_{ind} = \frac{\beta_{ind\_mt} + \beta_{ind\_pt}}{2} = \frac{2+\alpha}{2+2\alpha}[-(1+\alpha)\beta_O + \beta_M + \beta_P]$$

Thus, by plug-in the unbiased estimators of $\hat{\beta}_O, \hat{\beta}_M, \hat{\beta}_P$ from marginal GWAS, we can get unbiased estimator for $\beta_{dir}, \beta_{ind\_mt}, \beta_{ind\_pt}$:

$$\hat{\beta}_{dir} = (2+\alpha)\hat{\beta}_O - \hat{\beta}_M - \hat{\beta}_P$$

$$\hat{\beta}_{ind\_mt} = -\left(1+\frac{\alpha}{2}\right)\hat{\beta}_O + \frac{3-\alpha^2}{2(1-\alpha^2)}\hat{\beta}_M + \frac{1-2\alpha-\alpha^2}{2(1-\alpha^2)}\hat{\beta}_P$$

$$\hat{\beta}_{ind\_pt} = -\left(1+\frac{\alpha}{2}\right)\hat{\beta}_O + \frac{1-2\alpha-\alpha^2}{2(1-\alpha^2)}\hat{\beta}_M + \frac{3-\alpha^2}{2(1-\alpha^2)}\hat{\beta}_P$$

$$\hat{\beta}_{ind} = \frac{2+\alpha}{2+2\alpha}[-(1+\alpha)\hat{\beta}_O + \hat{\beta}_M + \hat{\beta}_P]$$

Several notes:

1) Genotype $(G_O, G_M, G_P)$ in these three GWASs can be from different trios
2) If at a SNP where mating is random, $\alpha = 0$, then the equations above become

$$\beta_{dir} = 2\beta_O - \beta_M - \beta_P$$

$$\beta_{ind\_mt} = -\beta_O + \frac{3}{2}\beta_M + \frac{1}{2}\beta_P$$

$$\beta_{ind\_pt} = -\beta_O + \frac{1}{2}\beta_M + \frac{3}{2}\beta_P$$

$$\beta_{ind} = -\beta_O + \beta_M + \beta_P$$

3) If there is no paternal effect, i.e. $\beta_{ind\_pt} = 0$, then we can rewrite the expression for $\beta_{dir}$ and $\beta_{ind\_mt}$:

$$\beta_{dir} = \frac{2}{3-\alpha^2}[(2+\alpha)\beta_O - (1+\alpha)\beta_M]$$

$$\beta_{ind\_mt} = \frac{2+\alpha}{3-\alpha^2}[-(1+\alpha)\beta_O - 2\beta_M]$$

4) If the maternal and paternal samples from different families are pooled together to run a parental GWAS (i.e., GWAS-MP) $Y_O = (Y_{OM}^T\ Y_{OP}^T)^T = G_{MP}\beta_{MP} + u_{MP} = (G_M^T\ G_P^T)^T\beta_{MP} + u_{MP}$, where $Y_{OM}$ and $Y_{OP}$ are the offspring's phenotypes whose

maternal genotype $G_M$ and paternal genotype $G_P$ are used in the GWAS. The parental effect size estimator

$$\hat{\beta}_{MP} = \frac{G_{MP}^T Y_O}{G_{MP}^T G_{MP}} = \frac{\sum_{i=1}^{n_M} G_M^T Y_{OM} + \sum_{i=1}^{n_P} G_P^T Y_{OP}}{\sum_{i=1}^{n_M} G_M^T G_M + \sum_{i=1}^{n_P} G_P^T G_P}$$

$$\to \frac{\left[n_M\left(\frac{1+\alpha}{2}\beta_{dir}+\beta_{ind\_mt}+\alpha\beta_{ind\_pt}\right)Var\left(G_{M,1}\right)+n_P\left(\frac{1+\alpha}{2}\beta_{dir}+\alpha\beta_{ind\_mt}+\beta_{ind\_pt}\right)Var\left(G_{P,1}\right)\right]}{n_M Var\left(G_{M,1}\right)+n_P Var\left(G_{P,1}\right)}$$

$$\to \frac{1+\alpha}{2}\beta_{dir} + \frac{n_M + \alpha n_P}{n_M + n_P}\beta_{ind\_mt} + \frac{\alpha n_M + n_P}{n_M + n_P}\beta_{ind\_pt}$$

Thus, we have

$$\beta_{MP} = \frac{1+\alpha}{2}\beta_{dir} + \frac{n_M + \alpha n_P}{n_M + n_P}\beta_{ind\_mt} + \frac{\alpha n_M + n_P}{n_M + n_P}\beta_{ind\_pt}$$
$$= \frac{n_M\beta_M + n_P\beta_P}{n_M + n_P},$$

which is a weighted effect size from $\beta_M$ and $\beta_P$.

a. If either $\beta_{ind\_mt} = \beta_{ind\_pt}$, or $n_M = n_P$

$$\beta_{MP} = \frac{1+\alpha}{2}\beta_{dir} + (1+\alpha)\beta_{ind},$$

then the direct and indirect effect sizes can be obtained using

$$\beta_{dir} = (2+\alpha)\beta_O - 2\beta_{MP}$$
$$\beta_{ind} = \frac{2+\alpha}{1+\alpha}\beta_{MP} - \frac{2+\alpha}{2}\beta_O$$

**Variances and covariances among effect size estimators**

Since the variances of $\hat{\beta}_{O,M,P}$ are reported in the GWAS summary statistics, the variances of $\hat{\beta}_{dir}$ and $\hat{\beta}_{ind}$ can be easily computed.

$$\mathrm{Var}\left(\hat{\beta}_{\mathrm{O,M,P}}\right) = \frac{\mathrm{Var}(Y_{\mathrm{O}})}{G_{\mathrm{O,M,P}}^T G_{\mathrm{O,M,P}}} \approx \frac{\mathrm{Var}(Y_{\mathrm{O}})}{n_{\mathrm{O,M,P}}\mathrm{Var}\left(G_{\mathrm{O,M,P}}\right)}$$

When there are overlapping sample among the input GWASs, their effect size estimates become correlated. Let's consider the overlapping case between GWAS-O and GWAS-M where a mother reported both her own and her offspring's phenotype $Y_{O,M}$, thus she was included in both GWAS-O and GWAS-M. Let's suppose the first $n_{OM}$ individuals in GWAS-O and GWAS-M are overlapping samples.

$$\begin{aligned}\mathrm{Cov}(\hat{\beta}_O, \hat{\beta}_M) &= \mathrm{Cov}\left(\frac{G_O^T Y_O}{G_O^T G_O}, \frac{G_M^T Y_M}{G_M^T G_M}\right) = \mathrm{Cov}\left(\beta_O + \frac{G_O^T u_O}{G_O^T G_O}, \beta_M + \frac{G_M^T u_M}{G_M^T G_M}\right) \\ &= E\left[\left(\beta_O + \frac{G_O^T u_O}{G_O^T G_O} - \beta_O\right)\left(\beta_M + \frac{G_M^T u_M}{G_M^T G_M} - \beta_M\right)\right] \\ &= E\left(\frac{G_O^T u_O}{G_O^T G_O}\frac{G_M^T u_M}{G_M^T G_M}\right) = \frac{G_O^T E(u_O u_M^T) G_M}{G_O^T G_O G_M^T G_M}\end{aligned}$$

$$\begin{aligned}E(u_o u_m^T) &= Cov(u_o, u_m^T) \approx Cov(Y_O, Y_M^T) = E(Y_O Y_M^T) \\ &= \begin{pmatrix} Cov(Y_{O,1}, Y_{M,1}) I_{n_{OM}\times n_{OM}} & 0 \\ 0 & 0\end{pmatrix} \in \mathbb{R}^{n_O \times n_M}\end{aligned}$$

Note, $u_{O,i}$ and $u_{M,j}$ are independent if either $i, j > n_{OM}$ or $i \neq j$. Since the effect from a single SNP is small, we used approximation $Y_{O,M} = G_{O,M}\beta_{o,M} + u_{O,M} \approx u_{O,M}$. Then,

$$\begin{aligned}\mathrm{Cov}(\hat{\beta}_O, \hat{\beta}_M) &\approx \frac{\sum_{i=1}^{n_{OM}} G_{O,i}^2 \,\mathrm{Cov}(Y_{O,1}, Y_{M,1})}{n_O n_M \mathrm{Var}(G_{O,1})\mathrm{Var}(G_{M,1})} \\ &\approx \frac{n_{OM}\mathrm{Cov}(Y_{O,1}, Y_{M,1})}{n_O n_M \mathrm{Var}(G_{O,1})} \\ &= \frac{n_{OM}\mathrm{Corr}(Y_{O,1}, Y_{M,1})\sqrt{\mathrm{Var}(Y_{O,1})\mathrm{Var}(Y_{M,1})}}{n_O n_M \mathrm{Var}(G_{O,1})} \\ &\approx \frac{n_{OM}\rho_{OM}}{\sqrt{n_O n_M}}\sqrt{\mathrm{Var}(\hat{\beta}_O)\mathrm{Var}(\hat{\beta}_M)}\end{aligned}$$

where $\rho_{OM} = \mathrm{Corr}(Y_{O,1}, Y_{M,1})$ is the phenotypic correlation among the overlapping samples. We used $Var(G_{O,i}) = Var(G_{M,i})$ in the second approximation since they are the variances of the same SNP in the overlapping sample. $\mathrm{Corr}(\hat{\beta}_O, \hat{\beta}_M) \approx \frac{n_{OM}\rho_{OM}}{\sqrt{n_O n_M}}$ can be estimated from the intercept of LDSC regression[1]:

$$E(z_{1j} z_{2j} | l_j) = \frac{\sqrt{n_O n_M}\rho_g}{M} l_j + \frac{n_{OM}\rho_{OM}}{\sqrt{n_O n_M}}$$

Similarly, if there are overlapping samples between GWAS-O and GWAS-P, we have

$$\mathrm{Cov}(\hat{\beta}_O, \hat{\beta}_P) \approx \frac{n_{OP}\rho_{OP}}{\sqrt{n_O n_P}}\sqrt{Var(\hat{\beta}_O)Var(\hat{\beta}_P)}$$

Since one cannot be a mother and father at the same time, there shouldn't be sample overlap between GWAS-M and GWAS-P. If there are real couples, however, $\hat{\beta}_M$ and $\hat{\beta}_P$ could be correlated due to assortative mating:

$$\mathrm{Cov}(\hat{\beta}_M, \hat{\beta}_P) \approx \frac{n_{MP}\rho_{MP}\alpha}{\sqrt{n_M n_P}}\sqrt{Var(\hat{\beta}_M)Var(\hat{\beta}_P)}$$

where $n_{MP}$ is the number of real couples and $\frac{n_{MP}\rho_{MP}\alpha}{\sqrt{n_M n_P}}$ is given by the LDSC intercept.

Then it's simple to derive the variances of the direct and indirect effect size estimates:

$$\begin{aligned}\mathrm{Var}(\hat{\beta}_{dir}) \quad &= (2+\alpha)^2\mathrm{Var}(\hat{\beta}_O) + \mathrm{Var}(\hat{\beta}_M) + \mathrm{Var}(\hat{\beta}_P) - 2(2+\alpha)\mathrm{Cov}(\hat{\beta}_O, \hat{\beta}_M) \\ &-2(2+\alpha)\mathrm{Cov}(\hat{\beta}_O, \hat{\beta}_P) + 2\mathrm{Cov}(\hat{\beta}_M, \hat{\beta}_P)\end{aligned}$$

$$\begin{aligned}\mathrm{Var}(\hat{\beta}_{ind\_mt}) \quad &= \left(1+\frac{\alpha}{2}\right)^2\mathrm{Var}(\hat{\beta}_O) + \left[\frac{3-\alpha^2}{2(1-\alpha^2)}\right]^2\mathrm{Var}(\hat{\beta}_M) + \left[\frac{1-2\alpha-\alpha^2}{2(1-\alpha^2)}\right]^2\mathrm{Var}(\hat{\beta}_P) \\ &- \frac{(3-\alpha^2)(2+\alpha)}{2(1-\alpha^2)}\mathrm{Cov}(\hat{\beta}_O, \hat{\beta}_M) - \frac{(1-2\alpha-\alpha^2)(2+\alpha)}{2(1-\alpha^2)}\mathrm{Cov}(\hat{\beta}_O, \hat{\beta}_P) \\ &+ \frac{(3-\alpha^2)(1-2\alpha-\alpha^2)}{2(1-\alpha^2)^2}\mathrm{Cov}(\hat{\beta}_M, \hat{\beta}_P)\end{aligned}$$

$$\begin{aligned}\mathrm{Var}(\hat{\beta}_{ind\_pt}) \quad &= \left(1+\frac{\alpha}{2}\right)^2\mathrm{Var}(\hat{\beta}_O) + \left[\frac{1-2\alpha-\alpha^2}{2(1-\alpha^2)}\right]^2\mathrm{Var}(\hat{\beta}_M) + \left[\frac{3-\alpha^2}{2(1-\alpha^2)}\right]^2\mathrm{Var}(\hat{\beta}_P) \\ &- \frac{(1-2\alpha-\alpha^2)(2+\alpha)}{2(1-\alpha^2)}\mathrm{Cov}(\hat{\beta}_O, \hat{\beta}_M) - \frac{(3-\alpha^2)(2+\alpha)}{2(1-\alpha^2)}\mathrm{Cov}(\hat{\beta}_O, \hat{\beta}_P) \\ &+ \frac{(3-\alpha^2)(1-2\alpha-\alpha^2)}{2(1-\alpha^2)^2}\mathrm{Cov}(\hat{\beta}_M, \hat{\beta}_P)\end{aligned}$$

$$\begin{aligned}\mathrm{Var}(\hat{\beta}_{ind}) = \left(\frac{2+\alpha}{2+2\alpha}\right)^2 &\left[\mathrm{Var}(\hat{\beta}_M) + \mathrm{Var}(\hat{\beta}_P) - (1+\alpha)^2\mathrm{Var}(\hat{\beta}_O) + 2\mathrm{Cov}(\hat{\beta}_M, \hat{\beta}_P)\right. \\ &\left. - 2(1+\alpha)\mathrm{Cov}(\hat{\beta}_M, \hat{\beta}_O) - 2(1+\alpha)\mathrm{Cov}(\hat{\beta}_P, \hat{\beta}_O)\right]\end{aligned}$$

**Effective sample sizes of the direct and indirect effects**

Many downstream analyses require sample sizes. Since the variance of the estimated effect size from a marginal regression is $\mathrm{Var}(\hat{\beta}_{O,M,P}) \approx \mathrm{Var}(Y_O)/n_{O,M,P}\mathrm{Var}(G_{O,M,P})$, we can compute an empirical effective sample size using

$$N_{dir,ind}^{eff} = \frac{\mathrm{Var}(Y_O)}{\mathrm{Var}(\hat{\beta}_{dir,ind})\mathrm{Var}(G_O)}$$

Note, in the above equation computing $\mathrm{Var}(G_O)$ requires minor allele frequency which may not be included in the GWAS summary statistics. Alternatively, we use $Var(\hat{\beta}_{O,M,P}) \approx \frac{Var(Y_O)}{n_{O,M,P}Var(G_{O,M,P})}$ in the formulas for $Var(\hat{\beta}_{dir,ind})$ derived in previous section to compute the effective sample sizes. This way, we don't need the minor allele frequency since it cancels out.

For case (i) in **Table 1** of the main text, we have

$$N_{dir}^{eff} = \left[\frac{(2+\alpha)^2}{n_o} + \frac{1}{n_M} + \frac{1}{n_P} - \frac{2(2+\alpha)l_{OM}}{\sqrt{n_O n_M}} - \frac{2(2+\alpha)l_{OP}}{\sqrt{n_O n_P}} + \frac{2l_{MP}}{\sqrt{n_M n_P}}\right]^{-1}$$

$$N_{ind}^{eff} = \left(\frac{2+2\alpha}{2+\alpha}\right)^2 \left[\frac{(1+\alpha)^2}{n_o} + \frac{1}{n_M} + \frac{1}{n_P} - \frac{2(1+\alpha)l_{OM}}{\sqrt{n_O n_M}} - \frac{2(1+\alpha)l_{OP}}{\sqrt{n_O n_P}} + \frac{2l_{MP}}{\sqrt{n_M n_P}}\right]^{-1}$$

where $l_{OM,OP,MP}$ are the correlations. When $\alpha = 0$ and no overlapping samples among input GWAS, their expressions can be simplified:

$$N_{dir}^{eff} = \left(\frac{4}{n_O} + \frac{1}{n_M} + \frac{1}{n_P}\right)^{-1}$$

$$N_{ind}^{eff} = \left(\frac{1}{n_O} + \frac{1}{n_M} + \frac{1}{n_P}\right)^{-1}$$

Similarly, for case (ii) in **Table 1**, the effective sample sizes are

$$N_{dir}^{eff} = \left[\frac{(2+\alpha)^2}{n_o} + \frac{4}{n_{MP}} - \frac{4(2+\alpha)l_{OMP}}{\sqrt{n_O n_{MP}}}\right]^{-1}$$

$$N_{ind}^{eff} = \left[\frac{(2+\alpha)^2}{4n_o} + \left(\frac{2+\alpha}{1+\alpha}\right)^2 \frac{1}{n_{MP}} - \frac{(2+\alpha)^2}{1+\alpha}\frac{l_{OMP}}{\sqrt{n_O n_{MP}}}\right]^{-1}$$

When $\alpha = 0$ and no sample overlap between the two input GWAS, we have

$$N_{dir}^{eff} = \left(\frac{4}{n_O} + \frac{4}{n_M}\right)^{-1}$$

$$N_{ind}^{eff} = \left(\frac{1}{n_O} + \frac{4}{n_M}\right)^{-1}$$

For case (iii) in **Table 1**, the effective sample sizes are

$$N_{dir}^{eff} = \left(\frac{3-\alpha^2}{2}\right)^2 \left[\frac{(2+\alpha)^2}{n_o} + \frac{(1+\alpha)^2}{n_M} - \frac{2(1+\alpha)(2+\alpha)l_{OM}}{\sqrt{n_O n_M}}\right]^{-1}$$

$$N_{ind}^{eff} = \left(\frac{3-\alpha^2}{2+\alpha}\right)^2 \left[\frac{(1+\alpha)^2}{4n_o} + \frac{1}{n_M} - \frac{(1+\alpha)l_{OM}}{\sqrt{n_O n_M}}\right]^{-1}$$

When $\alpha = 0$ and no overlapping samples, we have

$$N_{dir}^{eff} = \frac{9}{4}\left(\frac{4}{n_O} + \frac{1}{n_M}\right)^{-1}$$

$$N_{ind}^{eff} = \frac{9}{4}\left(\frac{1}{4n_O} + \frac{1}{n_M}\right)^{-1}$$

Note when deriving the effective sample sizes, we assumed $Var(G_O) = Var(G_M) = Var(G_P)$. This is because $N_{dir,ind}^{eff}$ are estimated from samples and in one GWAS multi-generational samples may exist or sample of a same generation may exist in different input GWAS.

### Effective sample sizes from a trio-based design

If using trios to run a full model $Y_O = G_O\beta_{dir} + G_M\beta_{ind\_mt} + G_P\beta_{ind\_pt} + \epsilon$, we can also compute the effective sample sizes for the direct and indirect effects. Let $G = (G_O, G_M, G_P)$ and $\beta = \left(\beta_{dir}, \beta_{ind\_mt}, \beta_{ind\_pt}\right)^T$. It can be found that (for simplicity we assumed $\alpha = 0$ here)

$$Var(\hat{\beta}) \approx (G^TG)^{-1}Var(Y_O) \approx \frac{Var(Y_O)}{Np(1-p)}\begin{pmatrix} 1 & -1/2 & -1/2 \\ -1/2 & 3/4 & 1/4 \\ -1/2 & 1/4 & 3/4 \end{pmatrix},$$

where $N$ is the number of trios (i.e., total $3N$ individuals), $p$ is the minor allele frequency. Using $N_{dir,ind}^{eff} = \frac{\mathrm{Var}(Y_O)}{\mathrm{Var}(\hat{\beta}_{dir,ind})2p(1-p)}$, we find the effective sample size are

$$N_{dir}^{eff} = \frac{N}{2}$$

$$N_{ind}^{eff} = N$$

Now let's compare them with those given by our framework. As were shown in the previous section, our effective sample sizes depend on assortative mating, sample overlap, and the phenotypic correlation $\rho$ (i.e., own vs. offspring phenotype) among overlapping samples. For simplicity, let's assume $\alpha = 0$ and equal number of total samples with sample overlap are used by our framework. That is, $n_O = n_M = n_P = N$ with $N/2$ overlapping samples between GWAS-O and GWAS-M, and the other $N/2$ GWAS-O samples are also in GWAS-P. Then, the correlation between effect size estimates $l_{OM} = l_{OP} = \rho/2$ and the effective sample sizes given by our approach are

$$N_{dir}^{eff} = \frac{N}{6 - 4\rho}$$

$$N_{ind}^{eff} = \frac{N}{3 - 2\rho}$$

where $\rho$ is the phenotypic correlation among the overlapping samples. When there's no sample overlap or $\rho = 0$, the effective sample sizes from the trio design are 3 times larger than those given by our study design. However, our approach becomes more and more powerful as $\rho$ approaches 1 (**Supplementary Figure 13**). Theoretically, our approach can be as powerful as the trio design if the phenotype correlation is 1.

When using GWAS-O and GWAS-MP as input (i.e., case ii of **Table 1**) with overlapping samples, i.e., $n_O = N, n_{MP} = 2N$ where all the GWAS-O samples are in GWAS-MP (i.e., this is when each individual reports both self and his/her offspring's phenotype, a study design that we are promoting in this paper), we will get the same effective sample sizes as shown above.

**Correlation between direct and indirect effect size estimates**

Our method introduces a technical correlation between the direct and indirect effect size estimates. For simplicity, let's consider the case of no sample overlap and random mating ($\alpha = 0$) among the input GWAS-O, GWAS-M, and GWAS-P:

$$\begin{aligned}\text{Cov}(\hat{\beta}_{dir}, \hat{\beta}_{ind}) &= \text{Cov}(2\hat{\beta}_O - \hat{\beta}_M - \hat{\beta}_P, -\hat{\beta}_O + \hat{\beta}_M + \hat{\beta}_P) \\ &= -2Var(\hat{\beta}_O) - Var(\hat{\beta}_M) - Var(\hat{\beta}_P) \\ &\approx -2\frac{Var(Y_{O1})}{n_O Var(G_{O1})} - \frac{Var(Y_{O2})}{n_M Var(G_{M2})} - \frac{Var(Y_{O3})}{n_P Var(G_{P3})} \\ &= -\frac{Var(Y_{O1})}{Var(G_{O1})}\left(\frac{2}{n_O} + \frac{1}{n_M} + \frac{1}{n_P}\right)\end{aligned}$$

In the above derivations, we assumed $Var(Y_O)$ and $Var(G)$ are the same among the three input GWASs. Their correlation is

$$Corr\left(\hat{\beta}_{dir}, \hat{\beta}_{ind}\right) = \frac{Cov\left(\hat{\beta}_{dir}, \hat{\beta}_{ind}\right)}{\sqrt{Var\left(\hat{\beta}_{dir}\right)Var\left(\hat{\beta}_{ind}\right)}} = -\frac{\frac{2}{n_O} + \frac{1}{n_M} + \frac{1}{n_P}}{\sqrt{\left(\frac{4}{n_O} + \frac{1}{n_M} + \frac{1}{n_P}\right)\left(\frac{1}{n_O} + \frac{1}{n_M} + \frac{1}{n_P}\right)}}$$

When $n_O = n_M = n_P$, $Corr\left(\hat{\beta}_{dir}, \hat{\beta}_{ind}\right) = -\frac{2}{3}\sqrt{2} \approx -0.94$. With increasing the sample sizes, the covariance between the estimated direct and indirect effects decreases. Their correlation, however, will not diminish since both their covariance and variances decrease at a same speed.

Similarly, when we use GWAS-O and GWAS-MP as inputs,

$$\begin{aligned}Cov(\hat{\beta}_{dir}, \hat{\beta}_{ind}) &= Cov(2\hat{\beta}_O - 2\hat{\beta}_M, 2\hat{\beta}_M - \hat{\beta}_O) \\ &\approx -\frac{Var(Y_{O1})}{Var(G_{O1})}\left(\frac{2}{n_O} + \frac{4}{n_M}\right)\end{aligned}$$

$$Corr(\hat{\beta}_{dir}, \hat{\beta}_{ind}) = \frac{Cov(\hat{\beta}_{dir}, \hat{\beta}_{ind})}{\sqrt{Var(\hat{\beta}_{dir})Var(\hat{\beta}_{ind})}} = -\frac{\frac{2}{n_O} + \frac{4}{n_M}}{\sqrt{(\frac{4}{n_O} + \frac{4}{n_M})(\frac{1}{n_O} + \frac{4}{n_M})}}$$

When $n_O = n_M$, $Corr\left(\hat{\beta}_{dir}, \hat{\beta}_{ind}\right) = -\frac{3}{10}\sqrt{10} \approx -0.95$.

### Using imputed parental genotypes

### Joint regression

Since genetic data of trios are limited, methods such as IMPISH[2] and SNIPar[3] were proposed to impute the parental genotypes using siblings. When only siblings' genotypes are available, these methods could impute the expected value of the sum of maternal and paternal genotypes. Define $G_C = G_M + G_P$ as the sum of parental genotypes.

Then the joint regression will give the estimates for the direct and indirect effects as we previously defined:

$$Y_O = \beta_{dir}^* G_O + \beta_C^* G_C + \epsilon_I$$

Let $X = (G_O, G_C)$, $\beta = (\beta_{dir}^*, \beta_C^*)^T$. The least square estimator for the model above is

$$\hat{\beta} = (X^T X)^{-1} X^T Y_O$$

Since

$$\begin{aligned}\left(\frac{1}{n}X^T X\right)^{-1} &= \begin{pmatrix} \frac{1}{n}G_O^T G_O & \frac{1}{n}G_O^T G_C \\ \frac{1}{n}G_C^T G_O & \frac{1}{n}G_C^T G_C \end{pmatrix}^{-1} \\ &\to \begin{pmatrix} 2p(1-p)(1+\alpha/2) & 2p(1-p)(1+\alpha) \\ 2p(1-p)(1+\alpha) & 4p(1-p)(1+\alpha) \end{pmatrix}^{-1} \\ &= \frac{1}{2p(1-p)(1+\alpha)} \begin{pmatrix} 2+2\alpha & -(1+\alpha) \\ -(1+\alpha) & 1+\alpha/2 \end{pmatrix}\end{aligned}$$

$$\begin{aligned}\left(\frac{1}{n}X^T Y\right) &= \frac{1}{n}\begin{pmatrix} G_O^T \\ G_C^T \end{pmatrix} \left(\beta_{dir} G_O + \beta_{ind\_\mathrm{mt}} G_M + \beta_{ind\_\mathrm{pt}} G_P + \epsilon\right) \\ &\to \begin{pmatrix} \beta_{dir} 2p(1-p)(1+\alpha/2) + (\beta_{ind\_mt} + \beta_{ind\_\mathrm{pt}}) p(1-p)(1+\alpha) \\ (\beta_{dir} + \beta_{ind\_\mathrm{mt}} + \beta_{ind\_\mathrm{pt}}) 2p(1-p)(1+\alpha) \end{pmatrix},\end{aligned}$$

$$\hat{\beta} \xrightarrow{d} \left(\beta_{\mathrm{dir}}, \frac{\beta_{\mathrm{ind\_mt}} + \beta_{\mathrm{ind\_pt}}}{2}\right)^T = (\beta_{\mathrm{dir}}, \beta_{\mathrm{ind}})^T$$

Therefore, in this joint regression, we get the unbiased estimates for the direct and indirect effect sizes:

$$\beta_{dir}^{*} = \beta_{dir}$$
$$\beta_{ind}^{*} = \beta_{ind}$$

**Marginal regression**

Alternatively, we could also run a marginal regression ($Y_O \sim G_C$), and then combined with the GWAS-O results to obtain the direct and indirect effects. From

$$Y_O = G_C\beta_C + u_C,$$

we find that

$$\hat{\beta}_C = (G_C^T G_C)^{-1} G_C^T Y_O \rightarrow \frac{Cov(G_C, Y_O)}{Var(G_C)}$$
$$= \frac{Cov\left(G_M + G_P, \beta_{dir}G_O + \beta_{ind\_mt}G_M + \beta_{ind\_pt}G_P + \epsilon\right)}{4p(1-p) + 4p(1-p)\alpha}$$
$$= \frac{1}{2}\beta_{dir} + \beta_{ind}$$

$$\beta_C = \frac{1}{2}\beta_{dir} + \beta_{ind}$$

Together with the expression for $\beta_O$ given in GWAS-O, we can get expressions for the direct and indirect effects

$$\beta_{dir} = (2+\alpha)\beta_O - 2(1+\alpha)\beta_C$$
$$\beta_{ind} = \frac{2+\alpha}{2}(2\beta_C - \beta_O)$$

Note the expressions for the direct and indirect effects are different from those obtained using GWAS-MP and GWAS-O. The reason is that in GWAS-MP we pooled parents to run linear regression but here we take the summation of parental genotypes. Thus, the correlation between fathers and mothers used in this GWAS is taken into consideration here. When $\alpha = 0$, they give exactly the same expressions.

### Polygenic score level direct and indirect effects dissection

#### Full model based on PGS

Our framework of dissecting direct and indirect effect on SNP level could be extended to polygenic score (PGS) level. Assume there are $L$ independent causal SNPs, then the polygenic model is

$$\begin{aligned}
Y_O &= \sum_{i=1}^{L}(\beta_{dir,i}G_{O,i} + \beta_{ind_{mt},i}G_{M,i} + \beta_{ind_{pt},i}G_{P,i}) + \epsilon \\
&= \sum_{i=1}^{L}\beta_{dir,i}\,(T_{M,i} + T_{P,i}) + \sum_{i=1}^{L}(\beta_{ind_{mt},i}T_{M,i} + \beta_{ind_{pt},i}T_{P,i}) \\
&+ \sum_{i=1}^{L}(\beta_{ind_{mt},i}NT_{M,i} + \beta_{ind_{pt},i}NT_{P,i}) + \epsilon \\
&= PGS_{\mathrm{dir}} + PGS_{ind,T} + PGS_{ind,NT} + \epsilon,
\end{aligned}$$

where we defined

$$PGS_{dir} = \sum_{i=1}^{L}\beta_{dir,i}\left(T_{M,i} + T_{P,i}\right) = \sum_{i=1}^{L}\beta_{dir,i}G_{O,i}$$

$$PGS_{ind,T} = \sum_{i=1}^{L}(\beta_{ind_{mt},i}T_{M,i} + \beta_{ind_{pt},i}T_{P,i})$$

$$PGS_{ind,NT} = \sum_{i=1}^{L}(\beta_{ind_{mt},i}NT_{M,i} + \beta_{ind_{pt},i}NT_{P,i})$$

are the PGS of transmitted alleles through direct pathway, the PGS of transmitted alleles through indirect pathway and the PGS of non-transmitted alleles through indirect pathway, respectively.

#### Joint PGS regression model

Consider the following regression model of direct and indirect PGS:

$$Y_O = \delta_{dir}\widetilde{\mathrm{PGS}}_{\mathrm{dir}} + \delta_{ind}\widetilde{\mathrm{PGS}}_{\mathrm{ind}} + \epsilon$$

where

$$PGS_{ind} = \sum_{i=1}^{L} \beta_{ind,i} \left(T_{M,i} + T_{P,i}\right) = \sum_{i=1}^{L} \beta_{ind,i}\, G_{O,i},$$

is the indirect PGSs using SNP level indirect effect sizes as the weights. $\widetilde{PGS}_{\mathrm{dir}}$ and $\widetilde{PGS}_{\mathrm{ind}}$ are their standardizations. Consider the least square estimators of the above model

$$\begin{pmatrix} \hat{\delta}_{dir} \\ \hat{\delta}_{ind} \end{pmatrix} = \left[\left(\widetilde{PGS}_{dir}, \widetilde{PGS}_{ind}\right)^T \left(\widetilde{PGS}_{dir}, \widetilde{PGS}_{ind}\right)\right]^{-1} \left(\widetilde{PGS}_{dir}, \widetilde{PGS}_{ind}\right)^T Y_O$$
$$= \begin{pmatrix} \widetilde{PGS}_{dir}^T \widetilde{PGS}_{dir} & \widetilde{PGS}_{dir}^T \widetilde{PGS}_{ind} \\ \widetilde{PGS}_{ind}^T \widetilde{PGS}_{dir} & \widetilde{PGS}_{ind}^T \widetilde{PGS}_{ind} \end{pmatrix}^{-1} \begin{pmatrix} \widetilde{PGS}_{dir}^T Y_O \\ \widetilde{PGS}_{ind}^T Y_O \end{pmatrix}.$$

Since $\widetilde{PGS}_{\mathrm{dir}}$ and $\widetilde{PGS}_{\mathrm{ind}}$ are standardized, $\widetilde{PGS}_{dir}^T \widetilde{PGS}_{dir} = \widetilde{PGS}_{ind}^T \widetilde{PGS}_{ind} = n$, where $n$ is the sample size. Denote that

$$\sigma_{11} = \sqrt{Var(PGS_{dir,1})} = \sqrt{\sum\nolimits_{i=1}^{L} \beta_{dir,i}^2 2p_i(1-p_i)(1+\alpha/2)},$$

$$\sigma_{22} = \sqrt{Var(PGS_{ind,1})} = \sqrt{\sum\nolimits_{i=1}^{L} \beta_{ind,i}^2 2p_i(1-p_i)(1+\alpha/2)},$$

$$\sigma_{12} = \sqrt{Cov(PGS_{dir,1}, PGS_{ind,1})} = \sqrt{\sum\nolimits_{i=1}^{L} \beta_{dir,i}\beta_{ind,i} 2p_i(1-p_i)(1+\alpha/2)},$$

Then,

$$\frac{1}{n} \widetilde{PGS}_{dir}^T \widetilde{PGS}_{ind} \to \frac{1}{\sigma_{11}\sigma_{22}} Cov(PGS_{dir,1}, PGS_{ind,1}) = \frac{\sigma_{12}^2}{\sigma_{11}\sigma_{22}}$$

and

$$n \begin{pmatrix} \widetilde{PGS}_{dir}^T \widetilde{PGS}_{dir} & \widetilde{PGS}_{dir}^T \widetilde{PGS}_{ind} \\ \widetilde{PGS}_{ind}^T \widetilde{PGS}_{dir} & \widetilde{PGS}_{ind}^T \widetilde{PGS}_{ind} \end{pmatrix}^{-1} \to \frac{1}{1 - \frac{\sigma_{12}^4}{\sigma_{11}^2 \sigma_{22}^2}} \begin{pmatrix} 1 & -\frac{\sigma_{12}^2}{\sigma_{11}\sigma_{22}} \\ -\frac{\sigma_{12}^2}{\sigma_{11}\sigma_{22}} & 1 \end{pmatrix}$$

Since

$$\begin{aligned}
\frac{1}{n}\widetilde{PGS}_{dir}^{T}Y_O &= \frac{1}{n\sigma_{11}}PGS_{dir}Y_O \\
&\to \frac{1}{\sigma_{11}}\sum_{i=1}^{L}\beta_{dir}^2 2\,p_i(1-p_i)(1+\alpha/2)+\beta_{dir}\beta_{ind}2p_i(1-p_i)(1+\alpha) \\
&= \frac{\sigma_{11}^2+\sigma_{12}^2+\sum_{i=1}^{L}\beta_{dir,i}\beta_{ind,i}2p_i(1-p_i)\,\alpha/2}{\sigma_{11}}
\end{aligned}$$

$$\begin{aligned}
\frac{1}{n}\widetilde{PGS}_{ind}^{T}Y_O &= \frac{1}{n\sigma_{22}}PGS_{ind}Y_O \\
&\to \frac{1}{\sigma_{22}}\sum_{i=1}^{L}\beta_{dir,i}\beta_{ind,i}2p_i(1-p_i)(1+\alpha/2)+\beta_{ind,i}^2 2p_i(1-p_i)(1+\alpha) \\
&= \frac{\sigma_{12}^2+\sigma_{22}^2+\sum_{i=1}^{L}\beta_{ind,i}^2 2p_i(1-p_i)\,\alpha/2}{\sigma_{22}}
\end{aligned}$$

Thus, estimated direct and indirect PGS effect sizes are

$$\begin{pmatrix}\hat{\delta}_{dir}\\ \hat{\delta}_{ind}\end{pmatrix}\to\begin{pmatrix}\sigma_{11}\\ \dfrac{1+\alpha}{1+\alpha/2}\sigma_{22}\end{pmatrix},$$

and

$$\delta_{dir}=\sigma_{11}=\sqrt{\sum\nolimits_{i=1}^{L}\beta_{dir,i}^2 2p_i(1-p_i)(1+\alpha/2)}$$

$$\delta_{ind}=\frac{1+\alpha}{1+\alpha/2}\sigma_{22}=\frac{1+\alpha}{1+\alpha/2}\sqrt{\sum\nolimits_{i=1}^{L}\beta_{ind,i}^2 2p_i(1-p_i)(1+\alpha/2)}$$

Therefore, if we regress the offspring's phenotype with direct and indirect PGSs (standardized) jointly in the model, their effect sizes give estimates for the effect sizes of the direct and indirect PGS. Note that the effect size for direct (indirect) PGS is only a function of the direct (indirect) effects $\beta_{dir,i}$ ($\beta_{ind,i}$) of all the causal SNPs, the minor allele frequency $p_i$ and assortative mating $\alpha$.

$\hat{\delta}_{dir}$ and $\hat{\delta}_{ind}$ can be readily obtained from direct and indirect effect summary statistics without using an external independent sample for regression. However, since there is $\beta_{dir,i}^2$ in the expression, estimators only using summary statistics would be very noisy. Running the multiple regression above with independent level data would give another estimation of the effect sizes which are independent with those from the summary statistics and hence reduce the noise.

**Marginal PGS regression models**

If we run a marginal PGS regression with either only direct or indirect PGS in the model, the regression coefficients are

$$\begin{aligned}
\delta_{dir}^* &= \left(\widetilde{PGS}_{dir}^T \widetilde{PGS}_{dir}\right)^{-1} \widetilde{PGS}_{dir}^T Y_O \\
&\to \frac{\sigma_{11}^2 + \frac{2+2\alpha}{2+\alpha}\sigma_{12}^2}{\sigma_{11}} = \frac{\sum_{i=1}^L \beta_{O,i}\beta_{dir,i} p_i(1-p_i)}{\sqrt{\sum_{i=1}^L \beta_{dir,i}^2 p_i(1-p_i)(1+\alpha/2)}} \\
&= \delta_{dir} + \frac{2+2\alpha}{2+\alpha}\frac{\sigma_{12}^2}{\sigma_{11}}
\end{aligned}$$

$$\begin{aligned}
\delta_{ind}^* &= \left(\widetilde{PGS}_{ind}^T \widetilde{PGS}_{ind}\right)^{-1} \widetilde{PGS}_{ind}^T Y_O \\
&\to \frac{\sigma_{12}^2 + \frac{2+2\alpha}{2+\alpha}\sigma_{22}^2}{\sigma_{22}} = \frac{\sum_{i=1}^L \beta_{O,i}\beta_{ind,i} p_i(1-p_i)}{\sqrt{\sum_{i=1}^L \beta_{ind,i}^2 p_i(1-p_i)(1+\alpha/2)}} \\
&= \delta_{ind} + \frac{2+2\alpha}{2+\alpha}\frac{\sigma_{12}^2}{\sigma_{22}}
\end{aligned}$$

Thus, the results of marginal regressions are not "pure" direct and indirect effect size but contain both the direct and indirect components.

**PGS level dissection**

We could extend the SNP level dissection idea presented in the main text to PGS level. Consider the following three separate regressions:

$$Y_O = \gamma_O \widetilde{PGS}_O + u_O,$$

$$Y_O = \gamma_M \widetilde{PGS}_M + u_M,$$

$$Y_O = \gamma_P \widetilde{PGS}_P + u_P,$$

where we regress the offspring phenotype $Y_O$ on the standardized own, maternal and paternal PGSs $\widetilde{PGS}_{\mathrm{O,M,P}}$, respectively. All PGSs are standardized. Denote

$$\sigma_O = \sqrt{Var(PGS_O)} = \sqrt{\sum_{i=1}^{L} \beta_{O,i}^2 2p_i(1-p_i)\left(1+\frac{\alpha}{2}\right)},$$

$$\sigma_M = \sigma_P = \sqrt{Var(PGS_M)} = \sqrt{Var(PGS_P)} = \sqrt{\sum_{i=1}^{L} \beta_{O,i}^2\, 2p_i(1-p_i)},$$

where the PGSs are calculated using the same weight $\beta_O$. Then the least square estimators in the marginal regressions

$$\hat{\gamma}_O = \frac{\widetilde{PGS}_O^T Y_O}{\widetilde{PGS}_O^T \widetilde{PGS}_O} \to \frac{1}{\sigma_O} Cov\left(PGS_{O,1}, Y_{O,1}\right)$$
$$= \frac{1}{\sigma_O} Cov\left(\sum_{i=1}^{L} \beta_{O,i} G_{O,i}\,, \sum_{i=1}^{L} \beta_{dir,i} G_{Oi} + \beta_{ind\_mt} G_{M,i} + \beta_{ind\_pt} G_{P,i} + \epsilon\right)$$
$$= \frac{1}{\sigma_O} \sum_{i=1}^{L} 2p_i(1-p_i)\beta_{O,i}\left[\beta_{dir,i}\left(1+\frac{\alpha}{2}\right) + \beta_{ind,i}(1+\alpha)\right]$$

$$\hat{\gamma}_M = \frac{\widetilde{PGS}_M^T Y_M}{\widetilde{PGS}_M^T \widetilde{PGS}_M} \to \frac{1}{\sigma_M} Cov\left(PGS_{M,1}, Y_{M,1}\right)$$
$$= \frac{1}{\sigma_M} Cov\left(\sum_{i=1}^{L} \beta_{O,i} G_{M,i}\,, \sum_{i=1}^{L} \beta_{dir,i} G_{Oi} + \beta_{ind_{mt}} G_{M,i} + \beta_{ind_{pt}} G_{P,i} + \epsilon\right)$$
$$= \frac{1}{\sigma_M} \sum_{i=1}^{L} p_i(1-p_i)\beta_{O,i}\left[\beta_{dir,i}(1+\alpha) + 2\beta_{ind_{mt},i} + 2\alpha\beta_{ind_{pt},i}\right]$$

$$\hat{\gamma}_P = \frac{\widetilde{PGS}_P^T Y_P}{\widetilde{PGS}_P^T \widetilde{PGS}_P} \rightarrow \frac{1}{\sigma_P} Cov\big(PGS_{P,1}, Y_{P,1}\big)$$

$$= \frac{1}{\sigma_P} Cov\left(\sum_{i=1}^{L} \beta_{O,i} G_{P,i}\,, \sum_{i=1}^{L} \beta_{dir,i} G_{Oi} + \beta_{ind\_mt} G_{M,i} + \beta_{ind\_pt} G_{P,i} + \epsilon\right)$$

$$= \frac{1}{\sigma_P} \sum_{i=1}^{L} p_i(1-p_i)\beta_{O,i}\left[\beta_{dir,i}(1+\alpha) + 2\alpha\beta_{ind\_mt,i} + 2\beta_{ind\_pt,i}\right]$$

Then effect sizes for the direct and indirect PGSs in marginal regressions could be obtained

$$2\,\hat{\gamma}_O - \frac{\sigma_M}{\sigma_O}(\hat{\gamma}_M + \hat{\gamma}_P) \rightarrow \frac{1}{\sigma_O} \sum_{i=1}^{L} 2\,\beta_{O,i}\beta_{dir,i} p_i(1-p_i) = \delta_{dir}^*,$$

$$\frac{(2+\alpha)\sigma_M}{(2+2\alpha)\sigma_O}(\hat{\gamma}_M + \hat{\gamma}_P) - \hat{\gamma}_O \rightarrow \frac{1}{\sigma_O} \sum_{i=1}^{L} 2\,\beta_{O,i}\beta_{ind,i} p_i(1-p_i) = \delta_{ind}^*$$

When $\alpha = 0$, they become

$$2\,\hat{\gamma}_O - (\hat{\gamma}_M + \hat{\gamma}_P) \rightarrow \frac{1}{\sigma_O} \sum_{i=1}^{L} 2\,\beta_{O,i}\beta_{dir,i} p_i(1-p_i) = \delta_{dir}^*,$$

$$(\hat{\gamma}_M + \hat{\gamma}_P) - \hat{\gamma}_O \rightarrow \frac{1}{\sigma_O} \sum_{i=1}^{L} 2\,\beta_{O,i}\beta_{ind,i} p_i(1-p_i) = \delta_{ind}^*,$$

which has the same form as those for the SNP level direct and indirect effects.
